# Tetrameric INTS6-SOSS1 complex facilitates DNA:RNA hybrid autoregulation at double-strand breaks

**DOI:** 10.1101/2024.02.19.580984

**Authors:** Qilin Long, Kamal Ajit, Katerina Sedova, Vojtech Haluza, Richard Stefl, Sadat Dokaneheifard, Felipe Beckedorff, Monica G Valencia, Marek Sebesta, Ramin Shiekhattar, Monika Gullerova

## Abstract

DNA double strand breaks (DSBs) represent a lethal form of DNA damage that can trigger cell death and initiate oncogenesis. The activity of RNA polymerase II (RNAPII) at the break site is required for efficient DSB repair. However, the regulatory mechanisms governing the transcription cycle at DSBs are not well understood. Here, we show that Integrator complex subunit 6 (INTS6) associates with the trimeric SOSS1 (comprising INTS3, INIP, and hSSB1) to form a tetrameric SOSS1 complex following DNA damage. INTS6 binds to DNA:RNA hybrids and plays a crucial role in Protein Phosphatase 2 (PP2A) recruitment to DSBs, facilitating the dephosphorylation of RNAPII. Furthermore, INTS6 prevents the accumulation of damage-induced RNA transcripts (DARTs) and the stabilization of DNA:RNA hybrids at DSB sites. INTS6 interacts with, and promotes the recruitment of Senataxin (SETX) to DSBs, facilitating the resolution of DNA:RNA hybrids/R-loops. Our results underscore the significance of the SOSS1 complex in the autoregulation of DNA:RNA dynamics and the promotion of efficient DNA repair.

## Introduction

The human genome is challenged by thousands of DNA lesions daily, originating from both endogenous and exogenous sources. Incorrect repair of DNA damage poses a threat to the genome stability (1). DNA double-strand breaks (DSBs) represent the most lethal form of DNA damage, involving the disruption of the structure of the DNA double helix (2). In eukaryotes, two major repair pathways, homologous recombination (HR) and non-homologous end joining (NHEJ) repair DSBs (1,3,4).

RNA polymerase II (RNAPII) is the enzyme, transcribing thousands of protein-coding genes and long non-coding RNAs across the genome. Unscheduled pausing of RNAPII may lead to the formation of R-loops, structures that consist of a DNA:RNA hybrid and a single strand DNA (ssDNA) stretch on the non-templated DNA strand (5). R-loops, usually found behind paused RNAPII, are generally considered to be a by-product of transcription and a potential threat to genome stability due to exposed single stranded DNA (6,7). Upon DNA damage, cells undergo transient global transcriptional repression, caused by physical blockage or degradation of RNAPII (8–10). Intriguingly, accumulating evidence illustrates that temporary damage-induced transcription activation at DSBs is required for efficient DNA repair (2,11–14). The *de novo* transcription at the DSBs leads to the production of nascent transcripts named damage-associated RNA transcripts (DARTs) or damage-induced lncRNAs (dilncRNAs), which serve as precursors for the generation of small DNA damage-derived RNAs (DDRNAs) (11,15,16). Although both DARTs and dilncRNAs are derived from DSBs, they possess distinct features. DARTs are strand specific transcripts generated by RNAPII phosphorylated on Y1 position (Y1P) at DSBs and can be characterised as primary DARTs (pri-DARTs), which are generated in direction away form DSBs, whist the secondary DARTs (se-DARTs), initiated from R-loops, are directed towards DSBs (11). In contrast, dilncRNAs (transcribed away from DSBs) are produced by RNAPII phosphorylated on S2 position (S2P) and S5 position (S5P), respectively. Specifically, the broken DNA ends, regardless of their genomic location, act as transcriptional promoters to form a pre-initiation complex, recruiting S2P and/or S5P modified RNAPII to produce dilncRNAs (15). Both DARTs and dilncRNAs are further processed by ribonuclease III enzymes Dicer and Drosha into DDRNAs, which facilitate efficient DNA repair through the recruitment of other DDR factors (11,15–18). DARTs and dilncRNA can form DNA:RNA hybrids at DSBs by annealing with the ssDNA overhang template after the end resection and R-loops by hybridising with ssDNA strand behind the paused RNAPII (18,19). DNA:RNA hybrids and R-loops are predominantly formed at DSBs in transcriptionally active loci (20–23) and can facilitate the regulation of repair pathway choice and the recruitment of repair factors such as BRCA1/2, RAD52, 53BP1, and RPA (11,19,23–26). However, the prolonged existence of R-loops near DSBs can lead to genome instability (21,27,28).

The Integrator is a multi-protein complex (>1.5 MD), which consists of at least 16 subunits (INTS1-15 and DSS/SEM1)(29), and has been reported to bind to and regulate RNAPII, modulating the transcription and RNA processing of various types of RNAs (29–32). It also participates in controlling RNAPII transcription initiation (33), pause release (33,34), elongation (34,35) and termination (36). However, the understanding of the functional roles of Integrator’s individual subunits or sub-complexes is very limited. Integrator subunits (INTS) 9 and INTS11, which share significant sequence conservation with the RNA endonucleases CPSF-100 and CPSF-73 respectively, exhibit similar functions in the cleavage of pre-mRNAs (29,32). Recently, INTS6, together with a noncanonical form of Protein Phosphatase 2A (PP2A) lacking the β subunit, has been shown to form an Integrator-PP2A complex, which is recruited to actively transcribing genes to oppose CDK9 kinase activity and to dephosphorylate RNAPII (37,38). PP2A is a dominant serine-threonine phosphatase, which participates in numerous cellular activities in various tissues. Nevertheless, its function in regulating the damage-induced transcription at DSBs remains elusive.

Notably, INTS3 and INTS6 play a role in DNA repair. INTS3, identified as a part of the heterotrimeric sensor of ssDNA (SOSS1) complex, along with hSSB1 (NABP2) and C9orf80 (INIP), contributes to efficient DNA repair(39). A similar complex consisting of INTS3 and INIP and hSSB2 (NABP1), named SOSS2 has also been implicated in DNA damage response (DDR) (40). INTS6 binds to the C-terminus of INTS3 *in vitro* (41), acting as a scaffold for hSSB1/2 and INIP, forming a tetrameric complex (42,43). However, the signals for the trimeric and tetrameric SOSS1/2 complexes assembly, coordination between subunits and their role in transcription at DSBs remain enigmatic. We recently showed that Abelson tyrosine kinase (c-Abl) phosphorylates hSSB1 as a part of the trimeric SOSS1 complex, which together with RNAPII, promotes the liquid-liquid phase separation at DSBs, enabling the formation of dynamic transient compartments for DNA repair(39). Another study highlighted a stable association between the SOSS1 complex and the Integrator-PP2A to facilitate promoter-proximal termination of RNAPII. The lack of SOSS1-Integrator-PP2A leads to increased RNAPII pausing and pervasive accumulation of R-loops leading to genome instability (44).

Senataxin (SETX) is a DNA:RNA helicase that resolves DNA:RNA hybrids in both damage and non-damage conditions (21,45,46). Mutations in SETX have been associated with multiple diseases, such as ataxia with oculomotor apraxia 2 or amyotrophic lateral sclerosis (47,48). Besides its roles in RNA processing, a recent study has revealed that SETX directly acts as a *bona-fide* RNAPII transcription termination factor (46). However, the precise mechanism governing the activity of SETX in DDR remains unclear.

In this study, we demonstrate that DNA damage stimulates the association of INTS6 with the trimeric SOSS1 to form the tetrameric complex, which subsequently recruits PP2A. INTS6 alone or as a part of tetrameric complex, binds to DNA:RNA hybrids and is required for the recruitment of PP2A to DSBs. Furthermore, INTS6 interacts with SETX and facilitates its localisation to damaged sites. Depletion of INTS6 resulted in increased levels of nascent DARTs/dilncRNAs and the accumulation of DNA:RNA hybrids at DSBs. Our data suggest that the co-ordinated activity of the INTS6-PP2A-SOSS1 complex and SETX drives the autoregulation of DNA:RNA hybrids at DSBs, promoting efficient DNA repair.

## Materials and Methods

### Plasmids

The ORF of INTS6 was cloned into plasmid 438C (pFastBac His6 MBP Asn10 TEV cloning vector with BioBrick Polypromoter LIC subcloning, Addgene plasmid #55220). Constructs 438B-INTS3, 438B-hSSB1, 438B-INIP, 438C-INTS6 were combined using BioBrick Polypromoter LIC subcloning into a single construct enabling co-expression of the four subunits of the tetrameric INTS6-SOSS1 complex from a single virus in insect cells. To generate plasmids enabling expression of the kinase module of TFIIH complex in insect cells, the ORFs for CDK7, MAT1, and CCNH were cloned into plasmid 438B and later combined into a single construct. Plasmid enabling expression of cABL^CAT^ (AA 83–534), alongside PTP1b^1-238^ was generously provided by Gabriele Fendrich and Michael Becker at the Novartis Institutes for Biomedical Research, Basel. Plasmid pGEX4T1-(CTD)_26_-(His)_7_ (provided by Olga Jasnovidova) was used to express and purify GST-(CTD)_26_-(His)_7_.

### Insect cell work

To generate viruses enabling the production of proteins in insect cells, the coding sequences and the necessary regulatory sequences of the constructs were transposed into bacmid using *E. coli* strain DH10bac. The viral particles were obtained by transfection of the bacmids into the Sf9 cells using FuGENE Transfection Reagent and further amplification. Proteins were expressed in 300 ml of Hi5 cells (infected at 1×10^6^ cells/ml) with the corresponding P1 virus at multiplicity of infection >1. The cells were harvested 48 hours post-infection, washed with 1x PBS, and stored at −80°C.

### Protein purification

#### Purification of MBP-INTS6 and INTS6-tetrameric SOSS complex

Pellets of Hi5 insect cells were resuspended in ice-cold lysis buffer [50 mM Tris pH 8.0; 500 mM NaCl; 0.4% Triton X-100; 10% (v/v) glycerol; 10 mM imidazole; 1 mM DTT; protease inhibitors (0.66 μg/ml pepstatin, 5 μg/ml benzamidine, 4.75 μg/ml leupeptin, 2 μg/ml aprotinin); and 25 U benzonase per ml of lysate]. The resuspended cells were gently shaken for 10 min at 4°C. To aid the lysis, cells were briefly sonicated. The cleared lysate was passed through 2 mL of Ni-NTA beads (Qiagen), equilibrated with buffer [50 mM Tris-HCl, pH 8; 500 mM NaCl; 10 mM imidazole; and 1 mM DTT]. Proteins were eluted with an elution buffer [50 mM Tris-HCl, pH 8; 500 mM NaCl; 1 mM DTT and 400 mM imidazole]. The elution fractions containing proteins were pooled, concentrated, and further fractioned on Superdex S-200 column (for MBP-INTS6 purification) or Superose 6 column (for INTS6-tetrameric complex purification) with SEC buffer [25 mM Tris-Cl pH7.5; 200 mM NaCl, 1 mM DTT]. Fractions were then concentrated, and glycerol was added to a final concentration of 10 % before they were snap-frozen in liquid nitrogen, and stored at −80 °C.

#### Purification of CTD polypeptides

Five grams of E. coli BL21 RIPL cells expressing GST-(CTD)_26_-(His)_7_ were resuspended in ice-cold lysis buffer [50 mM Tris-HCl, pH 8; 0.5 M NaCl; 10 mM imidazole; 1 mM DTT], containing protease inhibitors (0.66 μg/ml pepstatin, 5 μg/ml benzamidine, 4.75 μg/ml leupeptin, 2 μg/ml aprotinin) at +4°C. Cells were opened up by sonication. The cleared lysate was passed through 2 mL of Ni-NTA beads (Qiagen), equilibrated with buffer [50 mM Tris-HCl, pH 8; 500 mM NaCl; 10 mM imidazole; and 1 mM DTT]. hSSB1 was eluted with an elution buffer [50 mM Tris-HCl, pH 8; 500 mM NaCl; 1 mM DTT and 400 mM imidazole]. The elution fractions containing hSSB1 were pooled, concentrated, and further fractioned on Superdex S-75 column with SEC buffer [25 mM Tris-Cl pH7.5; 200 mM NaCl, 1 mM DTT]. Fractions containing pure hSSB1 were concentrated, glycerol was added to a final concentration of 10 % before they were snap-frozen in liquid nitrogen, and stored at −80 °C.

### Electrophoretic Mobility-Shift Assay (EMSA)

Increasing concentrations of the tested proteins (22, 44, 88, 167 nM) were incubated with fluorescently labelled nucleic acid substrates (final concentration 10 nM) in buffer D [25 mM Tris-Cl, pH 7.5, 1 mM DTT, 5 mM MgCl2 and 100 mM NaCl] for 20 min at 37°C. Loading buffer [60 % glycerol in 0.001% Orange-G] was added to the reaction mixtures and the samples were loaded onto a 7.5 % (w/v) polyacrylamide native gel in 0.5 x TBE buffer and run at 75 V for 1h at +4°C. The different nucleic acid species were visualised using an FLA-9000 Starion scanner and quantified in the MultiGauge software (Fujifilm). To calculate the relative amount of bound nucleic acid substrate the background signal from the control sample (without protein) was subtracted using the *band intensity - background* option. Nucleic acid-binding affinity graphs were generated with Prism-GraphPad 7.

### *In vitro* pull-down experiments

Purified GST, GST-CTD, GST-Y1P-CTD, or GST-S5,7P-CTD (5 μg each), respectively, were incubated with the SOSS complex (tetrameric) (5 μg) in 30 μl of buffer T [20 mM Tris-Cl, 200 mM NaCl, 10 % glycerol, 1 mM DTT, 0.5 mM EDTA, and 0.01% Nonidet P-40; pH 7.5] for 30 min at 4°C in the presence of GSH-beads. After washing the beads twice with 100 μl of buffer T, the bound proteins were eluted with 30 μl of 4xSDS loading dye. The input, supernatant, and eluate, 7 μl each, were analysed on SDS-PAGE gel.

### Micro-scale thermophoresis (MST)

Binding affinity comparisons via microscale thermophoresis were performed using the Monolith NT.115 instrument (NanoTemper Technologies). The CTD polypeptides (CTD, Y1P-CTD, and S5,7P CTD, respectively) were fused with msfGFP and served as ligands in the assays. Affinity measurements were performed in the MST buffer [25 mM Tris-Cl buffer, pH 7.5; 150 mM NaCl; 1 mM DTT; 5% glycerol; and 0.01% Tween-20]. Samples were soaked into standard capillaries (NanoTemper Technologies). Measurements were performed at 25°C, 50% LED, medium IR-laser power (laser on times were set at 3 s before MST (20 s), and 1 s after), constant concentration of the labelled ligand (20 nM), and increasing concentration of the trimeric SOSS complex (4.8–1200 nM, CTD-GFP and Y1P-CTD-GFP; 28.7-7250 nM, S5,7P CTD) or the tetrameric SOSS complex (3.4-846 nM, CTD-GFP and Y1P-CTD-GFP; 4.3-607.5 nM, S5,7P CTD), respectively. The data were fitted with Specific binding Hill Slope in GraphPad Prism software.

### Cell lines and cell culture

Cells were maintained in high-glucose DMEM medium (Life Technologies, 31966047) with 10% (vol/vol) fetal bovine serum (Merck, F9665-500mL), 2 mM L-glutamine (Life Technologies, 25030024) and 100 units/ml penicillin-streptomycin solution (Life Technologies, 15140122) at 37 °C with 5% CO_2_ supplement. The frequent mycoplasma test was conducted, and regular cell morphology authorization was performed with microscope. HeLa wild-type (WT) cells were obtained from ATCC. Wild-type U2OS or *AsiSI*-ER U2OS cells are gifts from the Legube Laboratory (CNRS – University of Toulouse, France). The stable INTS6-GFP and hSSB1-GFP mutants were generated in HeLa WT with Lipofectamine LTX (Invitrogen, 15338100) transfection (2μg plasmid) followed by 500 μg/ml hygromycin B (Gibco, 10687010) selection for 10 days before being single-cell sorted. Monoclonals were progressively grown until sufficient confluency before use. The colony with the highest GFP signal was further validated with western blot.

### Drugs and antibodies

Ionising Radiation (IR)-induced DNA damage was performed by using the CS-137 source (Gravatom, GRAVITRON RX30/55). Generally, IR=10Gy, and samples were harvested 10 min post-IR unless stated differently. Cells were incubated with 20μM triptolide (TPL) (Enzo life science, BV-1761-1) for 1h or 100μM 5,6-Dichloro-1-beta-D-ribofuranosylbenzimidazole (DRB) (Cayman Chemical, 10010302) for 2h or 1μM THZ1 (Stratech Scientific, A8882-APE-10mM) for 2h prior to the induction of DNA damage. 2.5μM LB-100 (Stratech Scientific, B4846-APE-5mg) was employed for 2h to inhibit PP2A. The break induction for wild-type U2OS or *AsiSI*-ER U2OS cells was achieved by using 400nM (Z)-4-hydroxy Tamoxifen (4-OHT) (Cayman Chemical, 14854-1mg-CAY) for 4h. The used antibodies were listed in Supplementary Table 4.

### RNA interference (RNAi) and plasmid transfection

RNA interference (25nM for siRAD51, 60nM for the rest siRNAs) was achieved with Lipofectamine RNAiMax (Life technologies, 13778075) in OPTI-MEM (Gibco, 11058021) by using reverse transfection method. The used siRNAs are listed in the Supplementary Table 4. The forward transfection method was used to deliver plasmids by using Lipofectamine 3000 (Invitrogen, L3000001) or Lipofectamine LTX (Invitrogen, 15338100). Gibson cloning was used to generate INTS6-GFP plasmid. DNA fragment (CDS) of INTS6 ordered from IDT, then inserted into pCMV3-C-GFPSpark® backbone (Sino Biological, HG22790-ACG) (PCR amplification with primers listed in Table S3) with NEBuilder HiFi DNA Assembly Cloning Kit (NEB, E5520S). The details of plasmids source and usage are listed in Supplementary Table 4. Plasmid sequence was authorised by sanger sequencing.

### Western blot

Cells after treatments were detached by trypsin (Life Technologies, 12604013) and resuspended with 1x Laemmli buffer [62.5 mM Tris pH6.8, 2% sodium dodecyl sulphate (SDS), 2% β-mercaptoethanol, 10% glycerol, 0.005% bromophenol blue] (Alfa Aesar, J61337AD) and boiled for 10min at 95 ⁰C before use. Sonication step (high power, 10s with probe sonicator) was included to reach a complete cell lysis. Each sample was centrifuge at full speed for 15min to pellet cell debris before loading onto a gel. NuPAGE™ 4 to 12%, Bis-Tris, 1.0 mm, Midi Protein Gels (Invitrogen™, WG1402BOX) (for BRCA1 detection) and 4–15% Mini-PROTEAN® TGX™ precast protein gels (BioRad, 4561083/4561086) (for the rest protein) were used with standard western blot process. Briefly, gel was electrophoresed in MOPS running buffer (for NuPAGE Bis-Tris gel) or Tris-Glycine running buffer (for PROTEAN® TGX™ gel) at 120V until proteins were separated. Subsequently, proteins on PROTEAN® TGX™ gel were transferred onto nitrocellulose membranes (Perkinelmer, NBA085A001EA) with Trans-Blot® Turbo™ Transfer System (Bio-Rad, 1704150) at 25V, constant 1.8A for 10min. For proteins on NuPAGE™ 4 to 12% Bis-Tris gel, wet transfer at 280mA for 2.5h at room temperature with icepack was employed. Membranes were then blocked with 5% non-fat milk (Sigma, 70166-500G) in PBS with 1% TWEEN-20 (Fisher Scientific, BP337-100) (PBST) at room temperature for 1 h. Primary antibodies listed in the Supplementary Table 4 was incubated overnight at 4⁰C. Proper secondary antibodies were used for 1h at room temperature before visualised the membranes with Pierce™ ECL Western Blotting Substrate (Thermo Scientific, 10005943) or SuperSignal™ West Pico PLUS Chemiluminescent Substrate (Thermo Scientific, 34580) and Amersham™ Hyperfilm™ ECL™ (VWR, 28-9068-35) film.

### Proximity Ligation Assay (PLA)

PLA was performed by using Duolink™ In Situ Red Starter Kit Mouse/Rabbit (Merck, DUO92101-1KT) according to the manufacturer’s instructions. 2 × 10^5^ cells were seeded onto glass coverslip (SLS, MIC3300) overnight before fixed with 4% paraformaldehyde (PFA) in PBS (Alfa Aesar, J61899) for 10min. After washing away 4%PFA with ice-cold PBS (5 times), permeabilization step was performed with 0.1% TritonX-100 (Merck, X100-100ML) in PBS for 10min before blocking (inside a humanity chamber) with 100μL blocking buffer from the kit for 1h at 37°C. The primary antibody was diluted to desire concentration with Duolink™ dilution buffer from the kit to desire concentration and incubate coverslips overnight at 4°C. Following primary antibody incubation, PLA probe incubation, ligation and amplification process were followed the manufacturer’s instructions. Coverslips were then mounted with DAPI solution from the kit and sealed onto clear slides and air-dry in dark before imaging with Olympus FluoView Spectral FV1200 confocal microscope with 60X oil immersion objective. Images were processed in FiJi software(49) quantified by using CellProfiler(50) 4.2.1 with sparkle counter pipeline.

Duolink™ In Situ Probemaker PLUS kit (Merck, DUO92009-1KT) was applied to conjugate PLA oligonucleotides (PLUS) to Y1P rat antibody for use in Duolink® PLA experiments by following the manufacturer’s protocol.

### Cell Lysis and Co-Immunoprecipitation (CoIP)

Approximately 1 × 10^7^ cells in a 15cm dish (at 50-70% confluency) were washed twice with PBS before being lifted by scrapping and collected via centrifuge (500g, 4⁰C, 5min) into a 1.5mL tube. The cell pellet was then lysed in 5X volumes of Lysis Buffer (300μL) [50mM Tris pH 8 (Merck, T6066), 150mM NaCl (Merck, S3014), 2.5mM MgCl_2_ (Merck, PHR2486), 1% NP40 (Merck, I8896-100ML), 10% Glycerol (Thermo, 032450.M1), 1X protease inhibitors (Merck, 11873580001)/1X phosphatase inhibitors (Thermo Fisher, A32961)(PPI)] and 1μL/sample of Benzonase® Nuclease (Merck, E1014-25KU) on wheel for 1h at 4°C with vigorous pipetting every 15min interval. The cell lysate was subsequently collected by full-speed centrifuge at 4⁰C for 10min before diluted with 1.5X cell volumes (450 μL) of dilution buffer [150mM NaCl, 2.5mM MgCl_2_, 10% Glycerol, 1X PPI] inside a fresh 2ml tube. 0.05-0.1X volume of diluted cell lysate was taken for Input. A mixture of Protein A agarose (Millipore, 16-157) and Protein G agarose (Millipore, 16-201) beads were blocked with 3%BSA in dilution buffer before adding to cell lysate for 1.5h at 4°C with rotation (50 μL/sample beads resuspended in 50 μL/sample dilution buffer) to achieve the preclearance. The pre-cleared cell lysate was incubated with antibody at 4°C overnight. The pulldown next day was performed with a mixture of 50 μL/sample Protein A+G agarose beads for 1.5h at 4⁰C. Then, the beads were wash with dilution buffer three times before eluted with 2X Laemmli Buffer and boiled for 10min at 95°C.

### Affinity purification of Flag-INTS6 or mock for mass spectroscopy

HEK293 stable cells overexpressing Flag-INTS6, or mock were cultured in DMEM media (Gibco, #11965-084) supplemented with puromycin and 10% FBS (Atlas Biologicals, #F-0500-D). The purification of nuclear Flag-INTS6 or mock was performed as described in Kirstein et al. (51). Briefly, the removal of the cytoplasmic fraction and nuclear lysates were extracted using 10 ml of the buffer containing 0.42 M NaCl, 20 mM Tris-HCl (pH 7.9), 1.5 mM MgCl2, 0.5 mM DTT, 25% glycerol, 0.2 mM EDTA, and 0.2 mM PMSF. The complex was purified using its incubation with 1 ml of anti-FLAG M2 affinity gel (Sigma) 6h at 4CC. Following spinning down the vial at 2000g for 2 min, pellet was washed twice with 10 ml of buffer BC500 (20 mM Tris (pH 7.6), 0.01% Triton X-100, 0.2 mM ETDA, 10 mM 2-mercaptoethanol, 10% glycerol, 0.2 mM PMSF and 0.5 M KCl), and four times with 10 ml of the buffer BC100 (0.01% Triton X-100, 20 mM Tris (pH 7.6), 10% glycerol, 0.2 mM EDTA, 100 mM KCl, 10 mM 2-mercaptoethanol, and 0.2 mM PMSF) and one final wash of BC100 without detergent. Following washing, affinity columns were eluted with 500ul of FLAG peptide solution (0.5ug/ul) resuspended in BC100 buffer.

#### Silver staining

The silver staining was performed as described in Kirstein et al. (51). Briefly, nuclear lysates of inputs and affinity-purified elution were loaded on a 4-20% Tris-glycine gel (Invitrogen, Cat# XP04205BOX), 10% acetic acid and 50% methanol was used to fix the gel in for 1 hour at room temperature. To complete fixation, the gel was transferred to 10% methanol and 7% acetic acid for 1 hour. For washing the gel, 10% glutaraldehyde was used for 15 min, followed by three times washing in MilliQ water for 15 min. Gel was stained for 15 min in 100 ml of staining solution (1g AgNO_3_, 2.8ml NH4OH, 185ul NaOH (stock 10N in MilliQ water), brought it up to 100 ml with MilliQ water). After washing 3 times for 2 min in MilliQ water, the gel was developed in 100 ml developing solution (0.5 ml 1% citric acid and 52 ul 37% formaldehyde in 100 ml MilliQ water). A solution containing 5% acetic acid and 50% methanol was used to stop the reaction.

#### Western blot

Nuclear lysates of inputs and affinity-purified elution were loaded onto 4-15 % Criterion TGX Stain-Free precast polyacrylamide gels (BIO-RAD, Cat# 5678085) and transferred to nitrocellulose membranes which were subsequently blocked by 5% BSA for 1 hour at RT. Membranes were incubated with primary antibodies (including: anti-INTS6 (R90 N-terminal homemade antibody), anti-INTS11 (Sigma Prestige, #HPA029025), anti-Senataxin (Abcam, 300439), anti-PP2A-C (CST, #2259), anti-PP2A-A (CST, #2041), and anti-RPB1 NTD (CST, #14958) for overnight at 4°C. Then following 3 times washing for 10 min, blots were incubated with HRP conjugated secondary antibodies for 30 min at RT. Western blot results were visualized and quantitated by iBright 1500 Imaging system.

### Chromatin Immunoprecipitation (ChIP)

ChIP and qPCR were performed by standard procedures as previously described(11). After the 4h DSB induction by 4OHT, 1 × 10^7^ wild-type U2OS or *Asi*SI-ER U2OS cells were crosslinked with 1% formaldehyde (Merck, 252549-100ML) for 10 min at 37°C and inactivated by the addition of glycine (Merck, G7126) to a final concentration of 125 mM for 10 min (at 37°C). Cells were detached with a cell lifter (Fisher Scientific, 11577692) and washed with ice-cold PBS twice (by spin 5min, 400g). The cell pellet was lysed with 500 μL of Cell Lysis Buffer [5 mM PIPES (VWR, 0169-100g), 85 mM KCl, 0.5% NP-40, 1X PPI] on ice for 10 min. The samples were then centrifuged at 800 rcf for 5min to remove cytoplasm fractions (supernatant). Nuclei pellet was resuspended with 400 μL Nuclear Lysis Buffer [50 mM Tris–HCl pH 8.0, 1% SDS (Merck,75746-1KG), 10 mM EDTA, 1X PPI] and incubated on ice for another 10 min. Following nuclei lysis, chromatins were sheared by Bioruptor® Pico (Diagenode) for 10min (medium power, 30s ON/OFF) to reach ∼500bp length. The supernatants containing the sheared chromatin were collected by centrifugation (14,000 rcf, 4°C, 10 min), and diluted with 2.5X volumes (∼1mL) Dilution Buffer [16.7 mM Tris-HCl pH8.0, 0.01 %SDS, 1.1% Triton X-100, 5mM EDTA, 167 mM NaCl, 1X PPI] and precleared by 30 μL protein A/G agarose beads (Merck-Millipore, 16-157/16-201) for 1h. 5μg of antibody was used to isolate protein-DNA complex overnight at 4°C (with rotation). The protein-DNA complex was pulled down by 40 μL protein A/G agarose beads for 1h at 4°C, and washed with Buffer A [20 mM Tris-HCl pH 8.0, 2 mM EDTA, 1% SDS, 0.1% Triton X-100 and 150mM NaCl] once, Buffer B [20 mM Tris-HCl pH 8.0, 2 mM EDTA, 0.1% SDS, 1% Triton X-100 and 500 mM NaCl] once, Buffer C [10 mM Tris-HCl pH 8.0, 1 mM EDTA, 1% NP-40, 1% Sodium Deoxycholate (D.O.C)(Merck, D6750-100G) and 250 mM LiCl (Merck, L4408-100g)] once and Buffer D [10 mM Tris-HCl pH 8.0 and 1 mM EDTA] twice. Then, protein-DNA complex was eluted from beads with Elution Buffer [1% SDS, 100 mM NaHCO_3_] by rotating at room temperature for 30min. To free DNA from protein-DNA complex, 55 μL Digest Buffer [400mM Tris-HCl pH6.5, 100 mM EDTA], 30 μL 5M NaCl (to reach 300mM), 1μL RNase A (10 μg/ml) (Thermo Scientific™, EN0531) and 2 μL Proteinase K (10 mg/ml) (Thermo Scientific™, EO0491) were added to sample tube and incubated at 65°C overnight. DNA was purified by phenol/chloroform (pH 7.0) (ThermoFisher, 10308293) and ethanol precipitation. qPCR was performed in triplicate using 1ng isolated genomic DNA in a 25 µl reaction containing SensiMix™ SYBR® (Scientific Laboratory Supplies, QT65005) and 10 µM each of forward and reverse primers (listed in Table S2) on Rotor-Gene Q (QIAGEN) with PCR procedures under the following programme: 1 cycle at 95°C for 10min; 45 cycles at 95°C, 15s and 62°C for 15s; 1 cycle at 72°C for 20s. The 2^ΔΔCt^ method was applied for quantification. Data are represented as mean ± SD.

### DNA:RNA hybrid immunoprecipitation (DRIP)

To preserve the native DNA:RNA hybrid structures, all the procedures were performed in cold room. The non-crosslinked AsiSI cells (5 × 10^6^ - 1 × 10^7^) were trypsinized and collected before incubated with 800 μL cell lysis buffer [85 mM KCl, 5 mM PIPES, 0.5% NP-40] for 10 min. After cell lysis, samples were spined at 500g, 4°C for 5min to pellet nuclei fractions and remove the cytoplasmic supernatant. The nuclei pellet was lysed in 800 μL nuclei lysis buffer [50 mM Tris-HCl pH 8.0, 5 mM EDTA, 1% SDS] for 10 min before subjected to proteinase K digestion (10 μL, 4h). Initially, 5 μL Proteinase K (10 mg/ml) (Thermo Scientific™, EO0491) were used and incubated at 55°C for 1h with vigorous pipetting every 15 minutes. After that, additional 3 μL Proteinase K was added to each tube for another 1h 55°C incubation. Another 2 μL Proteinase K was added to each tube for additional 2h incubation (55°C). Chromatin were subsequently precipitated by 5M KAc and isopropanol, washed with 75% Ethanol. After the air-dry of chromatin, 100 μL DEPC H_2_O was used to resuspend chromatin at room temperature for 3min and diluted with 300 μL IP Dilution Buffer [16.7 mM Tris-HCl pH8.0, 0.01 %SDS, 1.1% Triton X-100, 1.2 mM EDTA, 167 mM NaCl, 1X PPI] before been sonicated by Bioruptor (Diagenode, middle power, 30s on, 30s off for 10 min). Samples were precleared with 25 μL/sample Protein A Dynabeads (Life technologies, 10002D) for 1h before incubated with 2.5 μg S9.6 antibody (sigma, MABE1095) for 10h at 4 °C. The pull-down of S9.6-DNA:RNA hybrids was achieved by 25 μL/sample Protein A Dynabeads for 1h. Then, DNA:RNA hybrids were washed by Buffer A [20 mM Tris-HCl pH 8.0, 2 mM EDTA, 0.1% SDS, 1% Triton X-100 and 150 mM NaCl] once, Buffer B [20 mM Tris-HCl pH 8.0, 2 mM EDTA, 0.1% SDS, 1% Triton X-100 and 500 mM NaCl] once, Buffer C [10 mM Tris-HCl pH 8.0, 1 mM EDTA, 1% NP-40, 1% Sodium Deoxycholate (D.O.C) and 250 mM LiCl] once and Buffer D [10 mM Tris-HCl pH 8.0 and 1 mM EDTA] twice. The elution of S9.6-DNA:RNA hybrids were achieved by rotating with 250 μL/sample elusion buffer [1% SDS, 100 mM NaHCO_3_] at room temperature for 30min twice. DNA:RNA hybrids were then free by 2h Proteinase K (55 °C) incubation. The standard phenol/chloroform (ThermoFisher, 10308293) process was used to extract DNA for the following qPCR. The detailed qPCR section is listed in the ChIP section.

### Chromatin associated RNA sequencing (ChrRNA-seq)

The wild-type U2OS or *Asi*SI-ER U2OS cells with proper RNAi reaching 50%-70% confluency in 15cm dish (∼4-8 million cells) before starting of work. For DSB induction, 400nM 4-OHT was added to culture medium for 4h. Cells were then washed and harvested by gently scrapping into 5mL PBS and pelleted by centrifuge (400g, 5min, 4°C). The cell pellets were lysed with 4 mL HLB+N buffer [10mM Tris-HCl (pH 7.5), 10 mM NaCl, 2.5 mM MgCl_2_, 0.5% NP-40], and underlaid with 1 mL HLB+NS buffer [10 mM Tris-HCl (pH 7.5), 10 mM NaCl, 2.5 mM MgCl_2_, 0.5% NP-40, 10% sucrose] before spin at 400g for 5min (4°C) to collect the nuclear pellets. Cytoplasmic fragment was collected for western blot to confirm the successful breakage of cells. Next, nuclear pellets were lysed and resuspended in 125 µL NUN1 buffer [20 mM Tris-HCl (pH 7.9), 75 mM NaCl, 0.5 mM EDTA, 50% glycerol], and chromatin was extracted by incubating in 1.2 mL NUN2 buffer [20 mM HEPES-KOH (pH 7.6), 300 mM NaCl, 0.2 mM EDTA, 7.5 mM MgCl_2_, 1% NP-40, 1M urea] on ice for 15min with interval vortex in every 3-4 min. The chromatin samples were then collected by full speed spin at 4°C for 15min. To digest DNA and chromatin-associated proteins, 2µL/sample Proteinase K(20ug/uL)(NEB,P8107S) and 1µL/sample Turbo DNase in 1X Turbo DNase Buffer (100µL/sample)(Thermo fisher, AM2238) was added to digest chromatin pellets at 37°C with 1000rpm shake until pellets were dissolved. RNA was extracted by Trizol/chloroform RNA extraction, and further cleaning (to get rid of DNA contamination) with Monarch® Total RNA Miniprep Kit (NEB, T2010S) by following manufacturer’s protocol. Sequencing libraries preparation was performed with the TruSeq Stranded Total RNA Sample Preparation Kit (Illumina) followed by paired-end sequencing on HiSeq2000 (Illumina).

### ChrRNA-Seq Data Processing

ChrRNA-Seq adapters were trimmed using Cutadapt (version 4.4) (https://cutadapt.readthedocs.io/en/stable/installation.html) in paired end mode and the quality of the resulting fastq files were assessed using FastQC (https://www.bioinformatics.babraham.ac.uk/projects/fastqc/). The trimmed reads were then aligned to human hg19 reference genome using STAR aligner (52). Each alignment file was then split using Samtools into two alignment files containing positively stranded and negatively stranded reads (https://www.htslib.org/)

### Metagene Plots

Strand specific coverage files containing CPM normalized read count per nucleotide position was generated for each alignment using deepTools bamCoverage (https://deeptools.readthedocs.io/en/develop/) (53). ComputeMatrix operation of deepTools was then performed on the strand separated bigwig files to calculate the CPM coverage in the 2.5kb flanking region of AsiSI site with bin size set at one. Bedtools intersect was used to find region of genes in 2.5kb flank of each annotated AsiSI site. Then custom python script was employed to annotate the bin values in the positively stranded matrix as sense or antisense based on whether they lie in same or opposite orientation of gene regions near AsiSI respectively. Similarly, the negatively stranded matrix was annotated as sense or antisense using the above logic. Only bins laying within gene regions were utilized for sense/antisense annotation. The sense matrices from positively and negatively stranded matrices were concatenated to form a combined sense matrix containing read coverage in sense orientation around AsiSI site. Antisense matrix was also crafted in the same manner to represent antisense matrix in 2.5kb flank region near each annotated AsiSI site. Antisense reads corresponding to different gene regions lying within same AsiSI site were summed to ensure that the matrix contains each AsiSI site as row with 5000 bins as columns representing antisense coverage in 2.5kb flank region of annotated AsiSI. Similar procedure was also followed to generate sense matrix. Matrix was then subdivided into different categories based on the known annotation of AsiSI sites as HR prone, NHEJ prone, uncut, highly transcriptionally active or transcriptionally less active sites. Sense and antisense matrix were then averaged across the AsiSI sites and plotted as line plots with separate scales using matplotlib python package. Fill plots representing read coverage in 2.5kb flank region of individual AsiSI sites were also created using matplotlib as replacement for IGV snapshots.

### PCA Plots

Bigwigsummary function of deeptools was employed in conjunction with plotPCA function to compare read coverage in 5kb region of BLESS 80 AsiSI sites between sample replicates.

### Box Plots

Coverage from sense and antisense matrices were used to make box plots representing CPM normalized reads in 500bp flank region of each AsiSI site. Coverage was calculated by summing CPM values in 500 bins centering DSB for each AsiSI site from sense and antisense matrix respectively. Box plots were then made using matplotlib python package and significance determined with 2 sample Wilcoxon Test from scipy python package. The comparison of CPM coverage between HR and NHEJ sites in 500bp flank region of AsiSI were performed using Mann Whitney U Test from scipy python package.

Fold changes across 500bp region flanking AsiSI were calculated by taking ratio of coverage in these regions between condition and control matrix for sense and antisense separately. Log2Fold Changes were then represented as box plots and significant difference in median values between sense and antisense were determined using 2-sample Wilcoxon test.

### Heatmaps

plotHeatmap function of deepTools was used to make the heatmaps with regions set as all annotated DSBs arranged in ascending order of cleavage efficiency. Separate heatmaps were generated using the above procedure for sense and antisense matrices.

### ChIP-Seq, DRIP-Seq data processing

SETX +4-OHT ChIP-Seq and S9.6 +4OHT DRIP samples were downloaded from Array Express (E-MTAB-6318). Read quality of the fastq files was checked using FastQC before and after adapter trimming. The trimmed reads were then mapped to hg19 genome following the standard ChIP-seq pipeline. The classic ChIP-seq pipeline consists of BWA (http://bio-bwa.sourceforge.net/) for alignment and samtools for duplicate removal (rmdup), sorting (sort) and indexing (index). Coverage files containing CPM normalized read count per nucleotide position was generated for each alignment file using deeptools bamCoverage (https://deeptools.readthedocs.io/en/develop/). ComputeMatrix operation of deepTools was then performed on the bigwig files to calculate the CPM coverage in the 2.5kb flanking region of AsiSI site with bin size set to one.

### BLESS Seq data processing

BLESS Seq (E-MTAB-5817) was processed using the same protocol as detailed in (54). Read count coverage was calculated for all annotated DSBs (+-500bp) using bedtools multicov. The sites were then ordered based on read count coverage for representing cleavage efficiency of DSB sites.

### Fluorescence-Activated Cell Sorting (FACS) for reporter assay

RNAi was performed and 24h later, 0.1 million of cells with corresponding siRNA were replated for another 24h. Next, the I-*Sce*I expression vector, pCBASceI plasmid (1.5 µg) (Addgene, 26477)(55), was transfected by using forward transfection with Lipofectamine 3000. After 48h, cells were washed by ice-cold PBS twice, trpsinised and collected in 400 µl 10%FBS in PBS on ice before running FACS. siBRCA1 works as positive control for HeLa HR reporter cells; the DNA-PK inhibitor Wortmannin (sigma, W3144-250UL) works as the positive control for HeLa NHEJ reporter cells. Briefly, after cell replacing for 6h, 1µM Wortmannin (Wort) was maintained in cell culture media until harvest. Sample data acquisition was achieved with CytoFLEX Flow cytometer (Beckman Coulter) and analyzed with FlowJo software.

### Clonogenic Assay

1000 cells with RNAi were reseeded into a 12-well plate for 24h before subjected to 0 and 2Gy IR. The plate was then return to grow at 37°C for 7-10 days until clear colonies formed. Colonies were fixed and stained with a buffer of 0.5% crystal violet (Sigma, C6158-100G) and 20% methanol (Merck, 32213-2.5L-M) for 1h before washing by ddH_2_O. Plates was scanned and quantified by ImageJ with ColoneArea plugin.

### Comet assay

5000 cells with RNAi (resuspended in 50µl PBS) were fixed and embedded in 50µl CometAssay LMAgarose (bio-techne, 4250-050-02) before being spotted on a Cometslide (bio-techne, 4250-050-03). At this point, the final concentration of low-melting gel is 0.5%. On-gel cell lysis was performed by placing the Cometslide into lysis buffer [2.5M NaCl, 0.1M EDTA, 10mM Tris-Base, 10% DMSO (freshly added), 1% TritonX-100 (freshly added), pH=10] at 4°C overnight. After rinsing with ddH_2_O, the Cometslide was immersed into running buffer [0.3M NaOH, 1mM EDTA, pH=13] to lose the chromatin for 1h at 4°C before being ran at constant 300mA for 0.5h. The neutralization step was performed with 0.4M Tris-base buffer (pH=7.5) for 5min at room temperature twice before the Cometslide been washed by 70% ethanol for 15min and air dried. The chromatin stained with 2μg/mL DAPI (BD Biosciences, 564907) in PBS for 5min followed by 5min ddH2O washing. Images was acquired with EVOS M7000 microscope with 10X objectives. ImageJ with OpenComet plugin was used to perform the tail moment quantification. The significance was determined by using unpaired Welch’s correction.

Details about all key reagents used in this study can be found in Supplementary Table 4. Details about sequences of oligonucleotides used in this study can be found in Supplementary Tables 1-3.

### Statistical Analysis

Statistical tests were performed in GraphPad Prism 9.4.1. All error bars represent mean ±CSD unless stated differently. Each experiment repeats at least 3 times (N=3). The Kolmogorov-Smirnov normality test was performed to test for a normal distribution. If data meets normal distribution, statistical testing was performed using the Student’s *t*-test, one-way ANOVA, or unpaired Welch’s correction (for comet assay analysis). If data did not show a normal distribution, Mann–Whitney test for two groups (non-parametric comparison for PLA foci analysis), or Dunn’s test with Bonferroni corrections for multiple group comparisons. Significances are listed as \**p*C≤C0.05, \*\**p*C≤C0.01, \*\*\**p*C≤C0.001, \*\*\*\**p*C≤C0.0001.

## Results

### INTS6 forms a tetrameric SOSS1 complex and binds to RNAPII in response to DNA damage

Previous research has demonstrated the *in vitro* formation of a tetrameric complex, involving Integrator complex subunit 6 (INTS6) and the trimeric SOSS1 complex, comprised of INTS3, hSSB1/2, and INIP (41–43). However, this interaction has not been validated *in vivo* nor within the context of DDR. Utilizing Proximity Ligation Assay (PLA), we observed a significant increase in the interaction between INTS3 and INTS6 following irradiation (IR) (**Supplementary Figure S1A**). Subsequently, we investigated whether knocking down INTS6 (**Supplementary Figure S1B**) influenced the assembly of the trimeric SOSS1 complex at DSBs. No reduction in PLA foci of INTS3/hSSB1 and γH2AX was detected in the absence of INTS6 (**Supplementary Figure S1C and D**), indicating that the recruitment of the trimeric SOSS1 complex to DSBs is not dependent on INTS6.

In a previous study, we showed that the trimeric SOSS1 complex interacts with RNAPII in response to DNA damage(39). Therefore, we tested whether the tetrameric SOSS1 complex played a role in the regulation of RNAPII transcription. Initially, we used the irreversible RNAPII inhibitor triptolide (TPL). Using PLA we observed a transcription-dependent recruitment of the tetrameric SOSS1 complex to DSBs, evidenced by the proximity of INTS3 or INTS6 to γH2AX with or without IR treatment in the presence or absence of triptolide (**Supplementary Figure S2A**). Subsequently, we explored the proximity of INTS6 to total RNAPII and active RNAPII phosphorylated at Ser2 (S2P), Ser5(S5P) and Tyr1(Y1P), respectively. We identified an interaction between INTS6 and all forms of RNAPII tested, with a significant increase following IR (**Supplementary Figure S2B**). Furthermore, inhibitors DRB (inhibitor of CDK9, phosphorylating Ser2 on the CTD) and THZ1 (inhibitor of CDK7, phosphorylating Ser5 on the CTD) significantly reduced the number of PLA foci between INTS6 and S2P- and S5P-RNAPII, respectively, which confirmed the specificity of these interactions (**Supplementary Figure S2C and D**).

To test whether INTS6 interacts directly with RNAPII, we conducted *in vitro* pull-down experiments using purified INTS6 and the tetrameric SOSS1 complex (**Supplementary Figure S3A and B**), and unphosphorylated GST-CTD, Y1P CTD, or S5,7P CTD, respectively. Our data indicate that all three tested variants of GST-CTD polypeptides are effectively pulled down by the tetrameric SOSS1 complex, but not by the isolated INTS6 alone, suggesting that formation of the complex is required for the direct interaction with RNAPII CTD (**Supplementary Figure S3C and D**). To provide a quantitative understanding of the binding between SOSS1 complexes and CTD polypeptides, we used microscale-thermophoresis (MST) analysis. The results revealed that both trimeric and tetrameric SOSS1 complexes exhibited similar affinities for unphosphorylated CTD and Y1P CTD (**Supplementary Figure S3E and F**). However, the tetrameric SOSS1 complex displayed higher affinity for S5,7P CTD (**Supplementary Figure S3G**) compared to the trimeric SOSS1 complex.

### INTS6 localizes to DSBs in DNA:RNA hybrid-dependent manner

A recent study has shown that the SOSS1-Integrator-PP2A complex binds to R-loops at the promoters of protein-coding genes through the hSSB1 subunit in non-damage conditions (44). Our previous work demonstrated that the trimeric SOSS1 complex, specifically through the hSSB1 subunit, binds to various nucleic acids (NA) structures, including ssDNA, DNA:RNA hybrids, and R-loops, whilst the INTS3 subunit alone did not exhibit binding to these NA substrates(39). INTS6 is a putative DEAD box helicase, which might bind to NAs. In this study, we utilized *in vitro* Electrophoretic Mobility Shift Assay (EMSA) to assess the binding properties of INTS6 to various NA substrates, including a 61-mer single-strand DNA (ssDNA), 21-mer ssDNA, DNA:RNA hybrids and R-loops. Our findings indicated that INTS6 binds to DNA:RNA hybrid and R-loops. Moreover, it binds ssDNA in length-dependent manner (**Figure 1A**). Similar binding pattern was observed for purified tetrameric SOSS1 complex (**Figure 1B**), suggesting that the inclusion of INTS6 into the tetrameric SOSS1 does not alter its NA binding properties.

**Figure 1.**
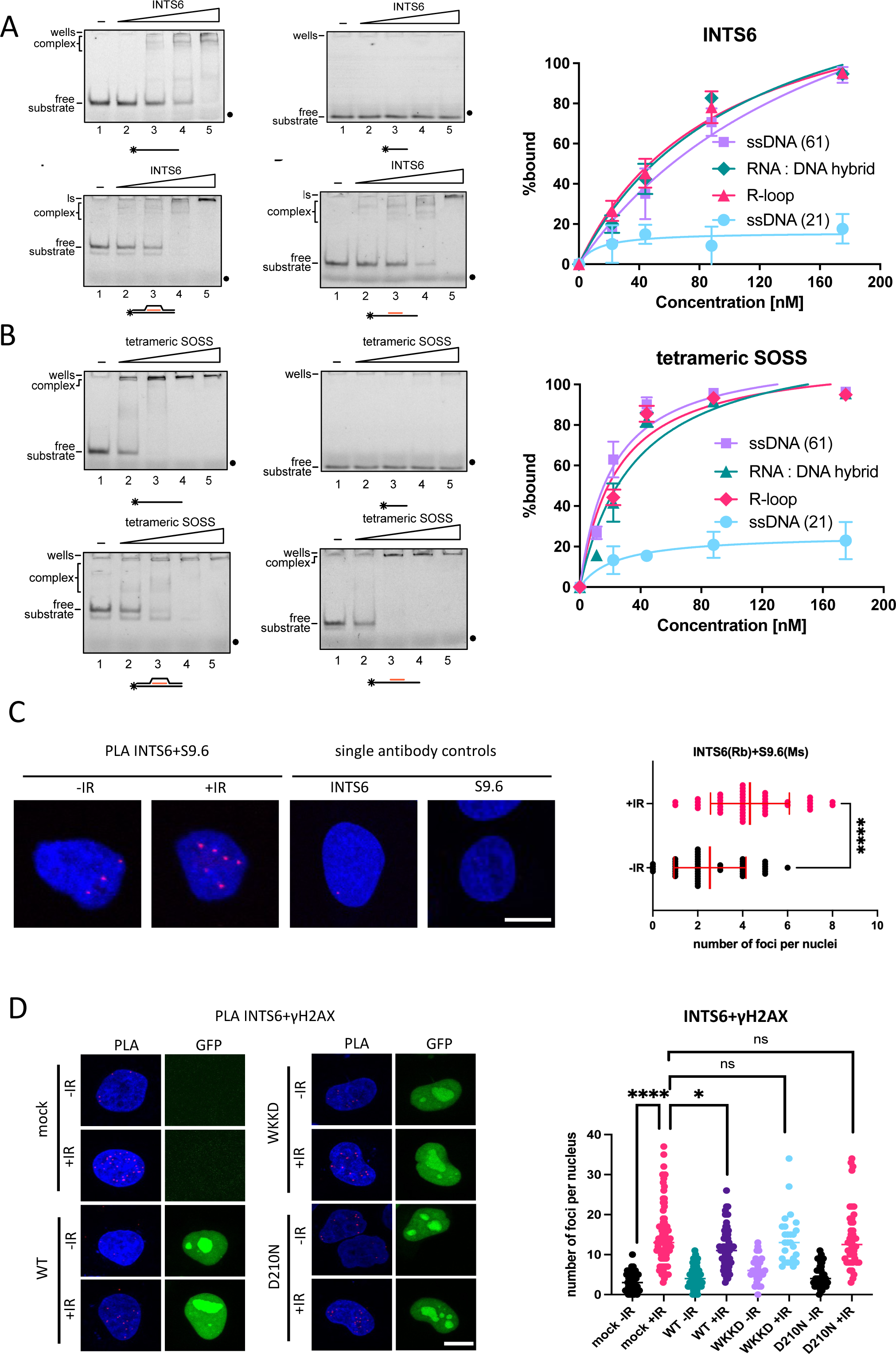
INTS6 localises to DSBs in DNA:RNA hybrid-dependent manner. **A)** Left: Scans of representative EMSA experiments of the INTS6 with 61- or 21-mer ssDNA, R-loops, and DNA:RNA hybrids. Right: Graph representing quantification of EMSA experiments (n=3). **B)** Left: Scans of representative EMSA experiments of the tetrameric SOSS1 complex with 61- or 21-mer ssDNA, R-loops, and DNA:RNA hybrids. Right: Graph representing quantification of EMSA experiments (n=3). **C)** PLA of INTS6 and S9.6 in cells with or without IR. Left: representative confocal microscopy images; right: quantification of left, error bar = mean ± SD, significance was determined using non-parametric Mann-Whitney test. ****p ≤ 0.0001. Scale bar =10μm. **D)** PLA of INTS6 and γH2AX in cells transiently transfected with RNAseH1^wt-GFP^ or RNAseH1^WKKD-GFP^ (binding and catalytic) or RNAseH1^D210N-GFP^ (catalytic) mutants with or without IR. Left: representative confocal microscopy images; right: quantification of left, error bar = mean ± SD, significance was determined using non-parametric Mann-Whitney test. ****p ≤ 0.0001, *p ≤ 0.05. Scale bar =10μm.

To test whether INTS6 is in close proximity to DNA:RNA hybrids in vivo, we performed a modified PLA using antibodies against INTS6 and S9.6 (recognizing DNA:RNA hybrids)(56) and observed a significant increased in PLA foci following IR (**Figure 1C**, single antibodies were used as a negative control).

Subsequently, we investigated whether R-loops or DNA:RNA hybrids play a role in the recruitment of INTS6 to DSBs *in vivo*. We transfected cells with plasmids expressing RNAseH1^WT-GFP^ (known to resolve DNA:RNA hybrids), RNAseH1^D210N-GFP^ (a catalytically inactive mutant), and RNAseH1^WKKD-GFP^ (a combined binding and catalytical mutant) (**Supplementary Figure S3H).** We then conducted PLA with antibodies against INTS6 and γH2AX. Our results revealed a significant increase in PLA foci in mock cells and cells expressing RNAseH1^D210N-GFP^ and RNAseH1^WKKD-GFP^, but not in those expressing RNAseH1^WT-GFP^. This suggests that INTS6 is recruited to DSBs in a DNA:RNA hybrid-dependent manner (**Figure 1D**).

### INTS6 is required for PP2A recruitment to DSBs

Integrator-PP2A complex plays a crucial role in dephosphorylating RNAPII at S2 and S5P of the CTD in non-damage condition. The phosphatase module formed by PP2A includes INTS6, and INTS6 aids in the assembly of PP2A into the Integrator-PP2A complex (38). Additionally, PP2A has been identified as binding to BRCA2 and promoting HR (57). Despite these known functions, the role of PP2A in transcription regulation at DSBs remains elusive.

To investigate whether the recruitment of PP2A to DSBs is dependent on INTS6, we employed PLA using antibodies against PP2A and γH2AX in control (mock) and INTS6 depleted cells in the presence or absence of IR. Our results revealed a significant increase in the interaction between PP2A and γH2AX upon IR, which was diminished in the absence of INTS6, indicating that INTS6 is necessary for the recruitment of PP2A to DSBs (**Figure 2A**). Subsequently, we explored the interaction between PP2A and RNAPII and detected an increased number of PP2A foci in response to DNA damage (**Supplementary Figure S4A**). Importantly, these PLA foci were significantly reduced upon INTS6 knockdown, emphasising the essential role of INTS6 in facilitating the binding of PP2A to RNAPII (**Supplementary Figure S4A**).

**Figure 2.**
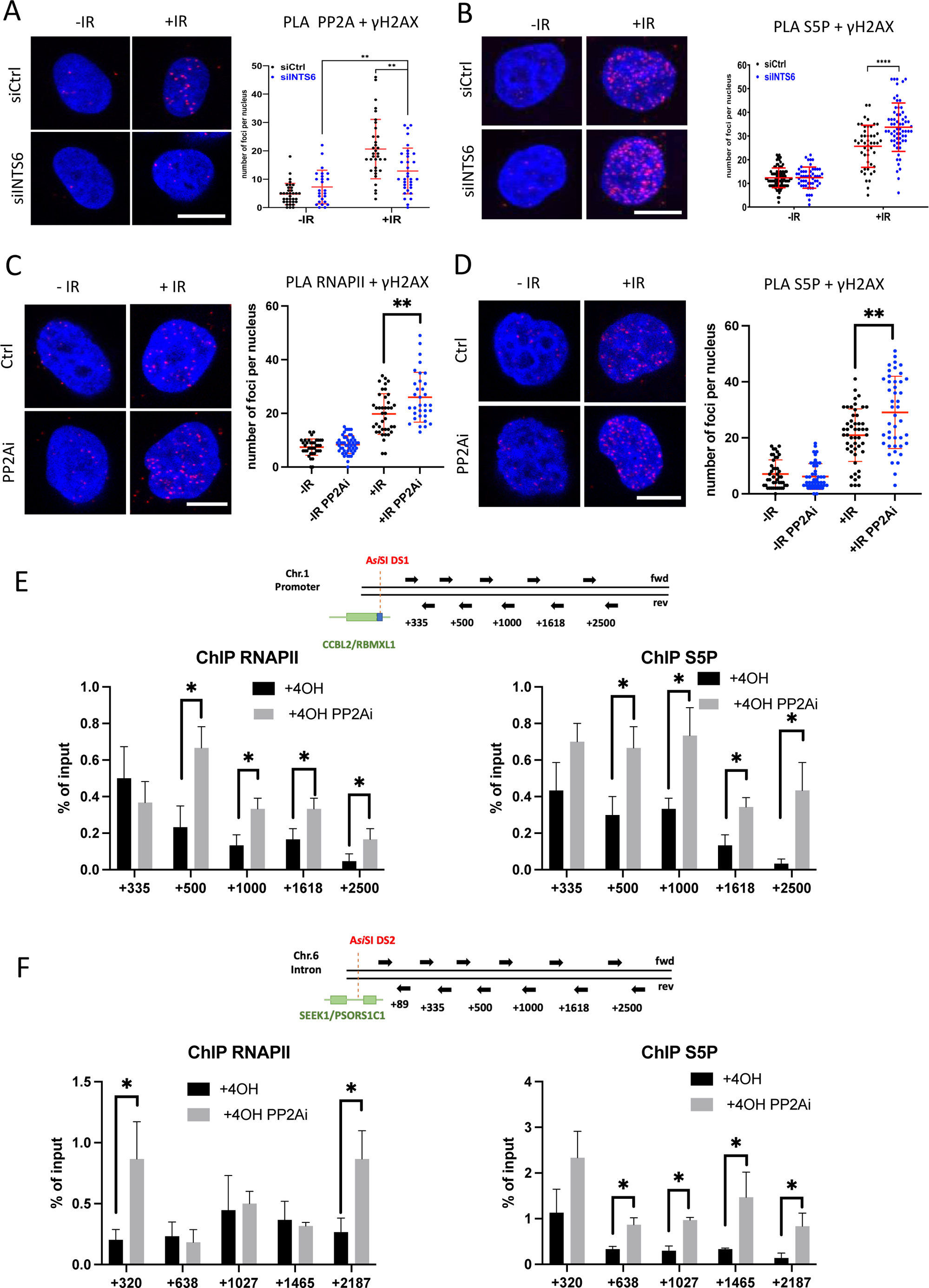
INTS6 facilitates PP2A recruitment to DSBs to dephosphorylate RNAPII. **A)** PLA of PP2A and γH2AX in wildtype or INTS6 knockdown cells with or without IR. IR=10Gy. Left: representative confocal microscopy images; right: quantification of left, error bar = mean ± SD, significance was determined using non-parametric Mann-Whitney test. **p ≤ 0.01. Scale bar =10μm. **B)** PLA of S5P and γH2AX in wildtype or INTS6 knockdown cells with or without IR. IR=10Gy. Left: representative confocal microscopy images; right: quantification of left, error bar = mean ± SD, significance was determined using non-parametric Mann-Whitney test. ****p ≤ 0.0001. Scale bar =10μm. **C)** PLA of RNAPII and γH2AX with or without IR in the presence or absence of PP2A inhibitor (LB-100, 2.5μM, 2h). IR=10Gy. Left: representative confocal microscopy images; right: quantification of left, error bar = mean ± SD, significance was determined using non-parametric Mann-Whitney test. **p ≤ 0.01. Scale bar =10μm. **D)** PLA as in C) for S5P and γH2AX **E)** Top: Drawing of ChIP probes positions around DS1. Bottom: bar charts showing RNAPII and S5P ChIP signals at DS1 in the absence or presence of PP2A inhibitor (LB-100, 2.5μM, 4h). n=3. Error bar = mean ± SD, significance was determined using Students t-test, unpaired *p ≤ 0.05 **F)** Top: Drawing of ChIP probes positions aournd DS2. Bottom: bar charts showing RNAPII and S5P ChIP signals over DS2 locus in the absence or presence of PP2A inhibitor (LB-100, 2.5μM, 4h). n=3. Error bar = mean ± SD, significance was determined using Students t-test, unpaired, *p ≤ 0.05.

In summary, these data suggest that INTS6 plays a critical role in facilitating the recruitment of PP2A to DSBs in a R-loop-dependent manner.

### PP2A is required for the dephosphorylation of RNAPII at DSBs

Next, we sought to determine whether PP2A plays a role in the dephosphorylation of RNAPII at DSBs. Initially, we observed an increased occupancy of active RNAPII phosphorylated at S5 at DSBs in cells depleted of INTS6, indicating impaired RNAPII dephosphorylation in the absence of INTS6. This defect may be attributed to the absence of PP2A at DSBs (**Figure 2B**). To further support this observation, we examined the occupancy of unmodified RNAPII, S5P RNAPII, and INT6 at DSBs induced by IR, following treatment with PP2A inhibitor LB-100. We detected significantly enriched PLA foci corresponding to the interaction between RNAPII, S5P RNAPII and γH2AX upon IR, and this effect was further intensified when PP2A was inhibited (**Figure 2C and D**). This suggests that PP2A inhibition may lead to the accumulation of phosphorylated RNAPII at DSBs. Interestingly, we also observed, after PP2A inhibition, significant increase in number of PLA foci corresponding to the proximity of INTS6 to γH2AX after PP2A inhibition (**Supplementary Figure S4B)**. As established earlier, INTS6 binds to phosphorylated RNAPII, and the increased number of INTS6/γH2AX PLA foci upon PP2A inhibition may be due to increased levels of S5P RNAPII at DSBs.

To study DSBs in a sequence-specific manner, we utilized the U2OS-A*si*SI-ER cell line, in which the A*si*SI restriction enzyme is fused to the oestrogen receptor (ER) ligand-binding domain. Addition of hydroxytamoxifen (4-OHT) induces translocation of the A*si*SI-ER enzyme to the nucleus, where it recognises 5′-GCGATCGC-3′ sequence motif and site-specific cuts simulating DSBs at specific genomic loci (58,59). There are 1231 predicted A*si*SI-ER cleavage sites in the human genome, but only 80 sites are efficiently cut *in vivo*, as it has been validated by γH2AX occupancy (12,58). Using this system, we showed that the Y1P RNAPII transcribes nascent RNAs (DARTs) (11). We selected two DSBs, DS1 (A*si*SI cut site in the promoter region of CCBL2/RBMXL1 gene on chromosome 1) and DS2 (in the intron 3 region of SEEK1/PSORS1C1 gene on chromosome 6) for further experiments and confirmed successful site-specific cuts by chromatin immunoprecipitation (ChIP) using γH2AX antibody (**Supplementary Figure S4C and D**). Subsequent ChIP experiments using RNAPII and S5P RNAPII antibodies revealed increased levels of RNAPII and S5P RNAPII around DS1 and DS2 upon PP2A inhibition (**Figure 2E and F**). The levels of RNAPII and S5P RNAPII were not affected at the GAPDH locus, used as a negative control (**Supplementary Figure S4E**).

The increased levels of S5P RNAPII at DSBs following PP2A inhibition, indicate that PP2A is required for the dephosphorylation of RNAPII at DSBs. Increased levels of RNAPII could reflect either the increased rate of transcription or impaired transcription termination.

### INTS6 depletion leads to the accumulation of damage-associated RNA transcripts (DARTs)

To investigate the impact of the accumulation of phosphorylated RNAPII in cells depleted of INTS6 on a genome-wide scale, we conducted chromatin-associated RNA-seq (chrRNA-seq) in U2OS-*Asi*SI-ER cells. Chromatin-associated RNAs were isolated from both control (siCtrl, scrambled siRNA) and INTS6-depleted cells (siINTS6) in the presence or absence of 4-OHT and subjected to next-generation sequencing (NGS). Principal component analysis revealed the clustering of sample replicates, showing robust data reproducibility **(Supplementary Figure S5A).**

Our analysis of chrRNA-seq revealed a significant increase in nascent transcripts surrounding DSBs (80 cut A*si*SI sites, called BLESS 80(54)) in INTS6 knockdown cells when compared to control samples (**Figure 3A**). In contrast, uncut sites (utilized as a negative control) exhibited consistent levels (**Figure 3B**). We categorized the nascent RNAs based on their direction relative to the transcriptional direction of genes at the AsiSI cuts, labelling them as sense (transcribed in the same direction as the nearby gene) or antisense (transcripts transcribed in the opposite direction as the nearby gene). Intriguingly, the depletion of INTS6 resulted in significantly increased levels of nascent RNAs in both directions, with a more pronounced increase in antisense RNA levels (**Figure 3A-D and Supplementary Figure S5B**). This observation was anticipated, as a certain amount of sense RNA is pre-synthesized before DSBs induction, whereas antisense RNA represents *de novo* transcription (**Figure 3E and F**).

**Figure 3.**
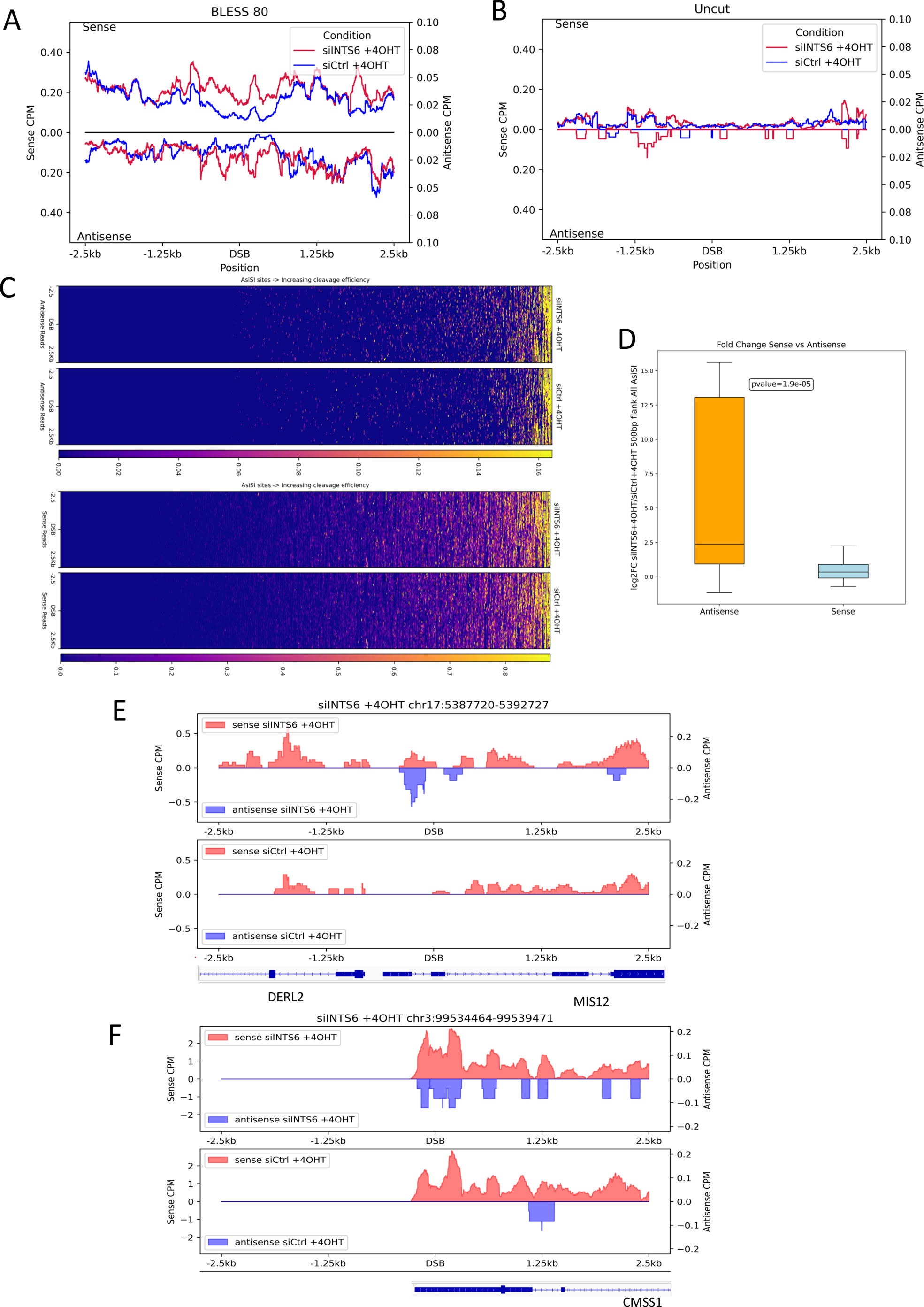
INTS6 depletion leads to the accumulation of Damage Associated RNA Transcripts (DARTs). **A)** Metagene plot shows chrRNA-Seq sense and antisense coverage in control (siCtrl) and INTS6 knockdown (siINTS6) cells with damage induction (+4-OHT) around 2.5kb flank region of BLESS 80 AsiSI sites (n=80). The reference genome is human hg19. **B)** Metagene plot shows chrRNA-Seq sense and antisense coverage in control (siCtrl) and INTS6 knockdown (siINTS6) cells with damage induction (+4-OHT) around 2.5kb flank region of uncut AsiSI sites (n=20). The reference genome is human hg19. **C)** Heatmaps show antisense (top) and sense (bottom) nascent RNA (chrRNA-seq) read coverage across all annotated AsiSI sites sorted based on cleavage efficiency in both INTS6 knockdown and control cells with damage induction (+4-OHT). **D)** Box plots show log2FoldChange of chrRNA-Seq coverage of sense reads and antisense reads upon INTS6 knockdown with damage induction compared to control with damage induction for all AsiSI (+/− 500bp). Wilcoxon 2 sample test is used for statistical testing of medians between sense and antisense log2FoldChange distribution. **E-F)** Representative snapshots of individual genes showing sense and antisense chrRNA-Seq coverage in INTS6 knockdown and control with damage induction around 2.5kb flank region of AsiSI cut. The specific loci information is listed on top of the snapshots. The reference genome is human hg19.

BLESS 80 cut sites can be characterised based on their proximity to highly transcribed regions(54). The analysis of chrRNA-seq in INTS6-depleted cells indicated a significant increase in both sense and antisense RNA transcripts around DSBs in proximity to highly transcribed genes, but only a significant increase in sense RNA transcripts near genes with low transcription activity (**Supplementary Figure S6A-D**). This supports the previous findings that pre-existed transcriptional states can influence DNA repair (60). Previous studies have shown that transcription and R-loops may act as a recruitment platform for HR repair factors (12). The *Asi*SI cut sites can be classified into HR or NHEJ prone DSBs based on the correlation ratio using ChIP-Seq data coverage of RAD51, HR factor, (*Asi*SI site +- 4kb) and XRCC4, known NHEJ factor, coverage (*Asi*SI site +- 1kb). The top 30 sites were annotated as HR prone due to a higher RAD51 coverage, and the bottom 30 as NHEJ prone due to a higher ratio of XRCC4(54). ChrRNA-seq analysis revealed a significant increase in nascent transcripts in INTS6-depleted cells for DSBs repaired by both pathways (**Supplementary Figure S6E-H**).

In conclusion, our data suggest that the depletion of INTS6 leads to elevated levels of nascent RNA around DSBs. This could imply that INTS6 either limits the rate of transcription at DSBs or facilitates the processing of DARTs.

### INTS6 interacts with Senataxin

The accumulation of DARTs in cells depleted of INTS6 suggests that INTS6, either directly or indirectly, plays a role in their processing. To identify proteins interacting with INTS6, we conducted affinity purification of nuclear Flag-INTS6. Silver stain of mock and Flag-INTS6 pull-downs indicated the presence of INTS6-specific bands, potentially corresponding to other Integrator subunits based on their molecular weight (**Figure 4A**). Indeed, mass spectrometry of Flag-INTS6 pull-downs confirmed the presence of all 15 subunits of the Integrator complex. Additionally, we identified subunits of RNAPII, PP2A kinase and SOSS complexes, including NABP1 (hSSB2), NAPBP2 (hSSB1) and INIP, in the INTS6 pull down samples. Notably, we also recovered Senataxin as an INTS6-associated polypeptide (**Figure 4B**). SETX, was described as an RNA/DNA helicase involved in R-loop resolution, transcription termination (45,46), DNA splicing, RNA processing, RNA stability (61) and coordination of replication and transcription conflicts (62). Additionally, SETX was recently identified to function as a *bona fide* transcription termination factor (46). We validated these findings by analysing our affinity-purified eluates of Flag mock and Flag-INTS6, with SETX, INTS11, RBP1, PP2A-A and PP2A-C specific antibodies. Specific bands for all tested proteins were observed in Flag-INTS6 samples, but not in Flag mock samples (**Figure 4C**).

**Figure 4.**
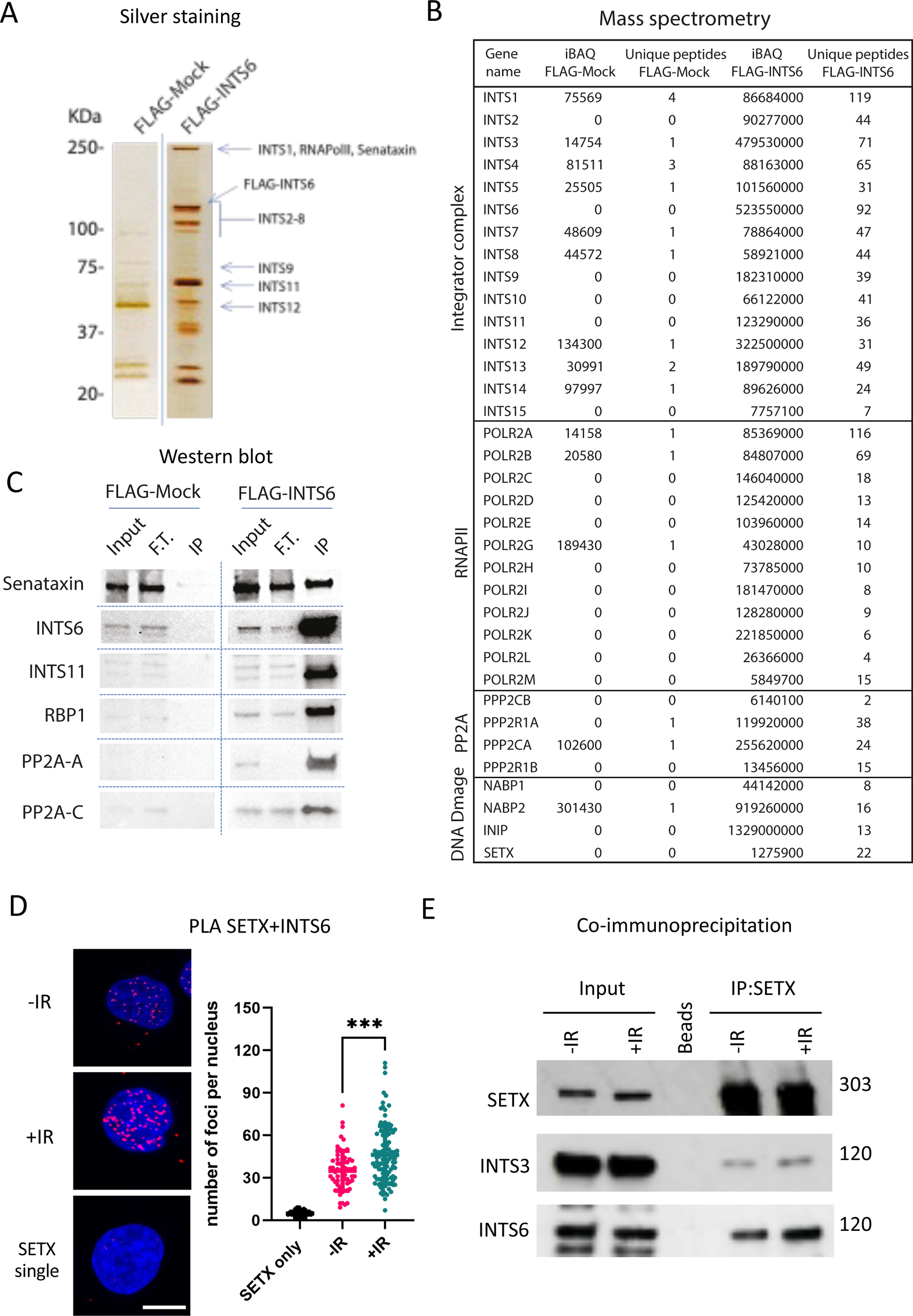
INTS6 associates with SETX. **A)** Silver stain of affinity-purified Integrator complex. The integrator complex was purified from nuclear lysate of HEK293Tcells, stably overexpressing Flag-INTS6. Mock indicates the same Flag-IP purification steps from parental HEK293T cells. The indicated Integrator subunits were assigned as identified by Baillat et al.(29). **B)** Affinity-purified Integrator complex mass spectrometry analyses were performed on nuclear lysate of HEK293Tcells stably overexpressing Flag-INTS6 or mock Flag-IP purification steps from parental HEK293T cells. The values represent intensity-based absolute quantification (iBAQ) intensities and the unique peptides. **C)** Affinity-purified Integrator complex followed by western blot indicated proteins. **D)** PLA of SETX and INTS6 with or without IR treatment. IR=10Gy. The single antibody was used as the negative control. Left: representative confocal microscopy images; right: quantification of left, error bar = mean ± SD, significance was determined using non-parametric Mann-Whitney test. ***p ≤ 0.001. Scale bar =10μm. **E)** Immunoprecipitation of SETX from cells with or without IR treatment, followed by Western blot showing signals for SETX, INTS6 and INTS3.

Subsequently, we investigated whether the INTS6-SETX interaction could be detected under DNA damage conditions. PLA revealed INTS6 in close proximity to SETX in non-damage condition, with this interaction further enhanced by DNA damage (**Figure 4D**). Finally, we repeated the co-IP using antibodies against endogenous SETX in cells with or without IR treatment. Western blot analysis confirmed the interaction between SETX and INTS6, with an augmentation of this interaction in DNA damage conditions (**Figure 4E**). Furthermore, we also detected INTS3 in SETX pull-down following IR, suggesting that DNA damage stimulates interaction between SETX and tetrameric SOSS1 complex.

In summary, these results revealed previously unreported interaction between INTS6 and SETX, which is stimulated by DNA damage.

### INTS6 is required for SETX recruitment and clearance of DNA:RNA hybrids at DSBs

We demonstrated that depletion of INTS6 leads to the accumulation of DARTs, particularly the secondary DARTs transcribed in the antisense direction towards DSBs. These secondary DARTs originate from R-loops formed behind paused RNAPII at the termination sites for primary DARTs(11). Consequently, we explored the potential requirement of INTS6 for the recruitment of SETX to DSBs. PLA confirmed the proximity of SETX to γH2AX upon IR treatment (**Figure 5A**). Interestingly, the interaction between SETX and γH2AX significantly decreased upon INTS6 knockdown (**Figure 5A**). Inhibition of PP2A (PP2Ai) enhanced SETX occupancy at DSBs (**Figure 5A**), suggesting that the accumulation of phosphorylated RNAPII may lead to failed RNAPII termination and increased stability of R-loops. This, in turn, could attract more SETX in a compensatory mechanism (**Figure 5A**). We further investigated the presence of SETX at specific DSBs by ChIP and observed significantly reduced levels of SETX at two selected DSBs in INTS6 knockdown samples. SETX was not detected at the control site (no DSB, chr22:23141639-23141780) or at the GAPDH locus (**Supplementary Figure S7A**). These results suggest an INTS6-dependent recruitment of SETX to the sites of DNA damage.

**Figure 5.**
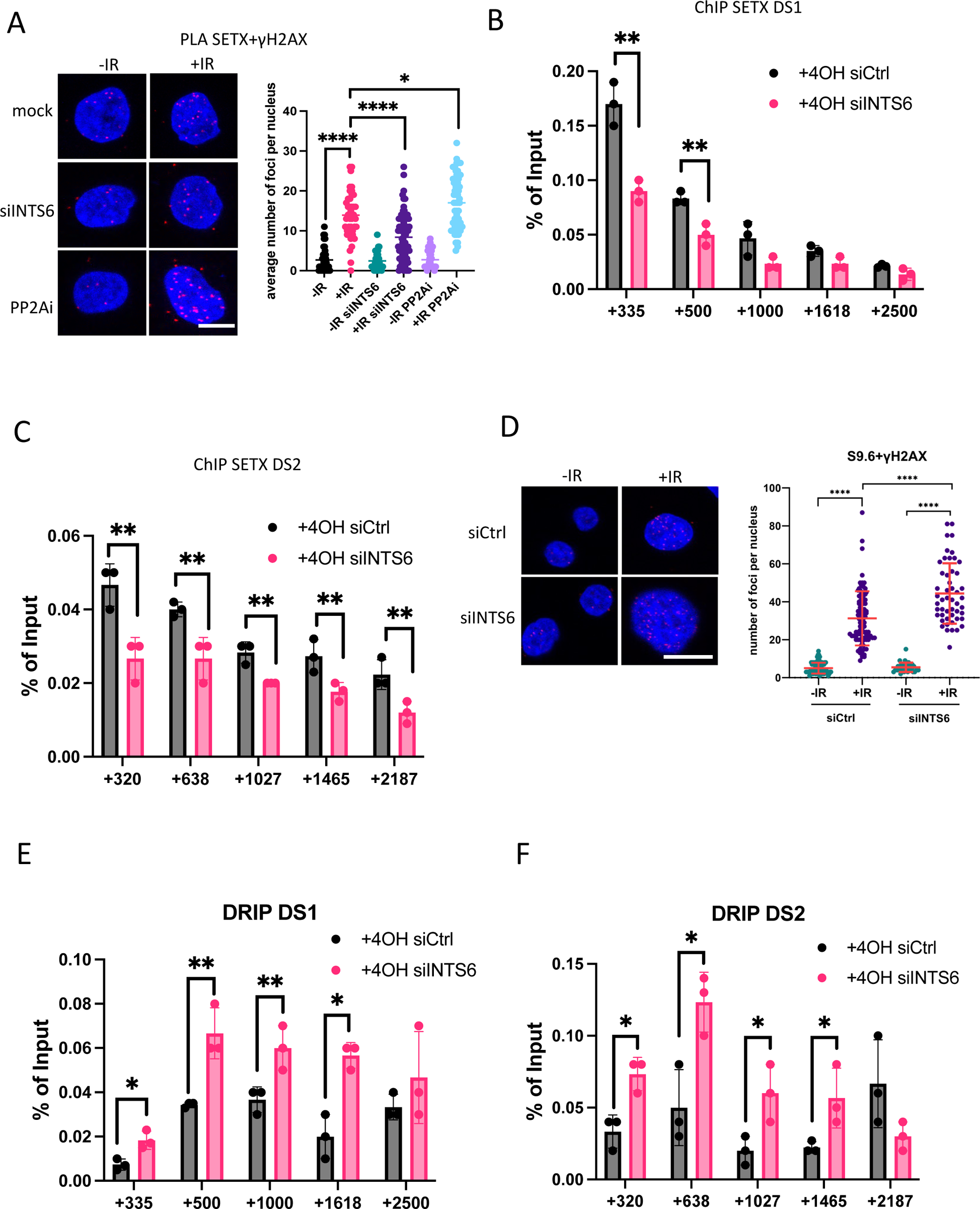
INTS6 is required for SETX recruitment to DSBs and clearance of DNA:RNA hybrids. **A)** PLA of SETX and γH2AX with or without IR in mock or siINTS6 cells or in the presence or absence of PP2A inhibitor (LB-100, 2.5μM, 2h). IR=10Gy. Left: representative confocal microscopy images; right: quantification of left, error bar = mean ± SD, significance was determined using non-parametric Mann-Whitney test. *p ≤ 0.05, ****p ≤ 0.0001. Scale bar =10μm. **B-C)** Bar charts showing SETX ChIP signals over DS1(B) and DS2(C) loci in the presence or absence of INTS6. n=3. Error bar = mean ± SD, significance was determined using Students’ t-test, unpaired, **p ≤ 0.01. **D)** PLA of S9.6 and γH2AX with or without IR in mock or siINTS6 cells. IR=10Gy. Top: representative confocal microscopy images; bottom: quantification of top, error bar = mean ± SD, significance was determined using non-parametric Mann-Whitney test. ****p ≤ 0.0001. Scale bar =10μm. **E-F)** Bar charts showing DRIP signals over DS1(E) and DS2(F) loci in the presence or absence of INTS6. n=3. Error bar = mean ± SD, significance was determined using Students’ t-test, unpaired, *p ≤ 0.05, **p ≤ 0.01.

SETX, functioning as a helicase that can resolve DNA:RNA hybrids at DSBs (21). To investigate whether SETX modulates levels of DNA:RNA hybrids in an INTS6-dependent manner upon DNA damage, we employed PLA in mock and INTS6-depleted cells exposed or not to IR treatment. We detected a significant increase in the levels of DNA:RNA hybrids levels upon IR, which was further increased by INTS6 depletion (**Figure 5D**). DNA:RNA immunoprecipitation (DRIP) analysis using the S9.6 antibody to capture DNA:RNA hybrid levels around DSBs, revealed increased levels of DNA:RNA hybrids in cells depleted of INTS6 at two selected DSBs. Levels of DNA: RNA hybrids at control loci: no DSB and GAPDH, remains unchanged (**Figure 5E and F, Supplementary Figure S7B**).

These findings suggest that the absence of INTS6 impairs the recruitment of SETX to DSBs, resulting in prolonged existence of DNA:RNA hybrids. This underscored the importance of INTS6 in maintaining the homeostasis of DNA:RNA hybrids at DSBs.

### INTS6-dependent accumulation of DARTs correlates with DNA:RNA hybrids at DSBs

DARTs and dilncRNA can form DNA:RNA hybrids and R-loops at DSBs by hybridising to the DNA overhangs after resection or unwound DNA behind paused RNAPII (11,15). Analyses of SETX ChIP-seq and S9.6 DRIP-seq in U2OS-A*si*SI-ER cells have indicated that SETX localizes to DSBs, playing a critical role in resolving DNA:RNA hybrids and safeguarding genome stability (21). In this study, we integrated chrRNA-seq datasets with SETX ChIP-seq and S9.6 DRIP-seq datasets to examine whether INTS6-dependent DARTs correlate with SETX and DNA:RNA hybrid occupancy around DSBs. Heatmaps displaying the levels of nascent RNA from control and siINTS6 chrRNA-seq data in conjunction with SETX ChIP-seq and S9.6 DRIP-seq datasets revealed a partial positive correlation between SETX and DARTs and negative correlation between SETX, DARTs, and DNA:RNA hybrids around DSBs (**Supplementary Figure 6A and Figure S8**). Box plot analysis of the nascent RNA transcript levels demonstrated that the INTS6-dependent increase in DARTs levels was most significant right at DSBs, decreasing with distance from DSBs (**Figure 6B**). Additionally, we generated a merged metagene plot showing nascent RNA levels, SETX-ChIP-seq and DRIP-seq for 80 cleaved *Asi*SI sites (**Figure 6C**) as well as for uncut sites (**Figure 6D**), using a 2.5kb frame size on both sides of DSBs. Interestingly, the metagene profile of SETX occupancy correlated with DARTs that were increased in an INTS6-dependent manner. Furthermore, the metagene profile of increased DARTs and SETX negatively correlated with the DRIP profile in control sample, suggesting that INTS6 depletion might lead to the accumulation DNA:RNA hybrids around DSBs (**Figure 6C-F**). Similar overlapping metagene patterns were also observed at both DSBs near sites with high and low transcriptional activity and HR and NHEJ-prone sites (**Supplementary Figure S9A-D**).

**Figure 6.**
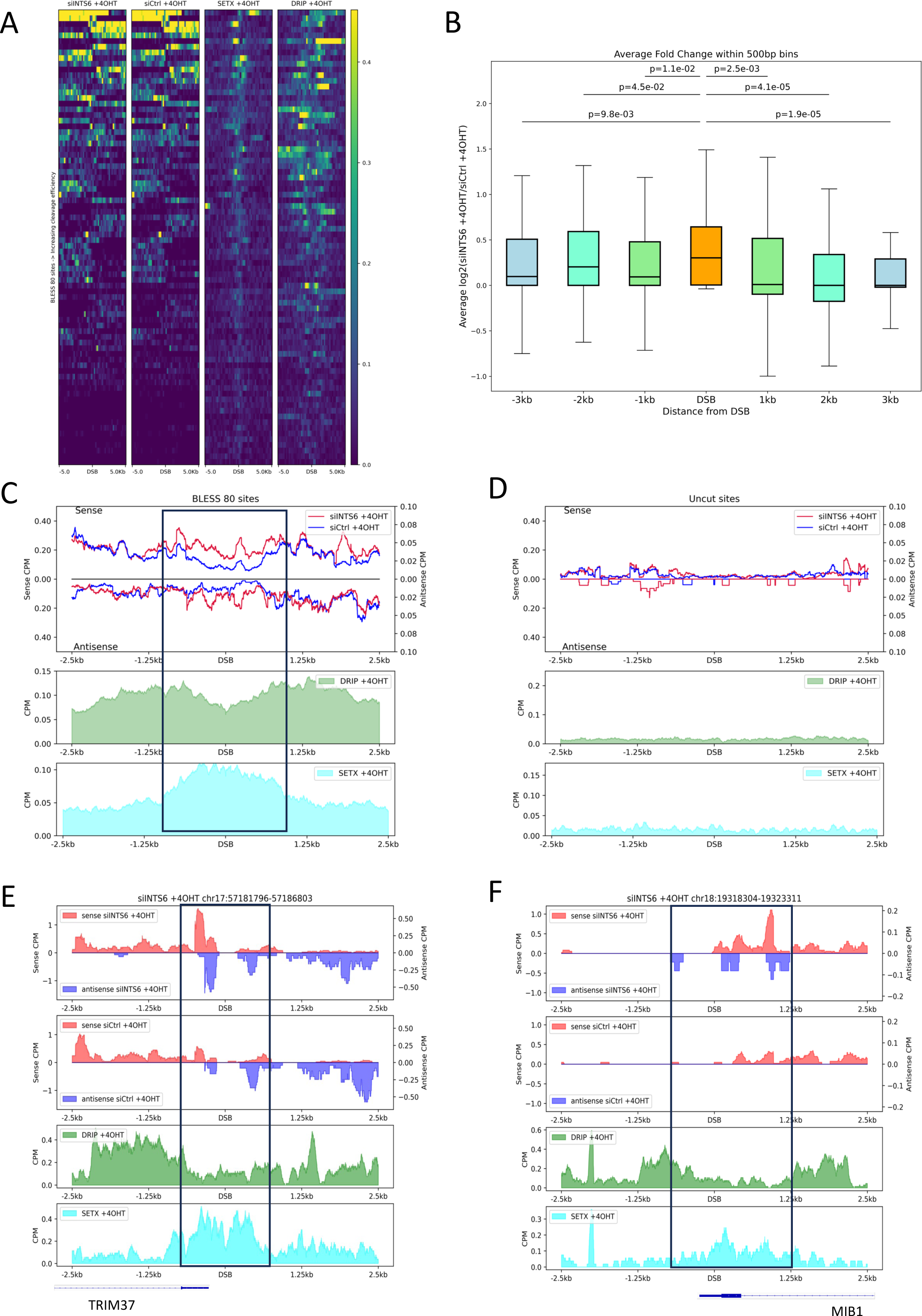
INTS6-dependent accumulation of DARTs correlates with DNA:RNA hybrids at DSBs. **A)** Heatmaps show the siINTS6 chrRNA-Seq coverage, Ctrl chrRNA-Seq coverage, SETX ChIP-Seq coverage and S9.6 DRIP-Seq coverage across BLESS 80 *Asi*SI sites sorted by cleavage efficiency during damage induction. The reference genome is human hg19. **B)** Box plots show log2FoldChange of total chrRNA-Seq coverage of INTS6 knockdown with damage induction compared to control with damage induction in 500bp bins centered at DSB, 1kb, 2kb and 3kb away from DSB (BLESS 80 *Asi*SI sites)(n=64, discard the outliners). The reference genome is human hg19. **C)** Metagene plots show S9.6 DRIP-Seq coverage and SETX coverage upon damage induction along with chrRNA-Seq sense and antisense coverage in siINTS6 and control with induced damage around 2.5kb flank region of BLESS 80 *Asi*SI sites (n=80). The reference genome is human hg19. **D)** Metagene plots show S9.6 DRIP-Seq coverage and SETX coverage upon damage induction along with chrRNA-Seq sense and antisense coverage in siINTS6 and control with induced damage around 2.5kb flank region of uncut *Asi*SI sites (n=20). The reference genome is human hg19. **E-F)** Representative snapshots of individual genes showing DRIP-Seq coverage and SETX coverage upon damage induction along with sense and antisense chrRNA-Seq coverage in siINTS6 and control with damage induction around 2.5kb flank region of *Asi*SI cut. The specific loci information is listed on top of the snapshot respectively. The reference genome is human hg19.

These findings support the notion that INTS6-dependent DARTs could contribute to DNA:RNA hybrids located at DSBs. INTS6, by binding to DNA:RNA hybrids, subsequently recruits SETX, which, in turn, resolves them. These results underscore the pivotal role of INTS6-mediated DNA:RNA hybrids autoregulation.

### INTS6 is required for efficient DNA damage repair

Intrigued by these findings, we sought to validate the significance of INTS6 in DDR. Initially, we conducted a clonogenic assay in control and INTS6 knockdown cells exposed to IR. The results revealed a growth defect attributed to INTS6 depletion, further exacerbated by IR treatment, leading to significant growth inhibition (**Figure 7A**). To investigate the specific DSB repair pathway in which INTS6 might be involved, we utilized reporter cell lines. The DR-GFP HR HeLa reporter cell contains a specially designed SceGFP sequence, with an I-*Sce*I cutting site, a stop codon and an in frame GFP template. Transient expression of the pCBA*Sce*I plasmid allows measurement of GFP-expressing cells by fluorescence-activated cell sorting (FACS) to assess HR repair efficiency. Depletion of INTS6 in this system resulted in robust HR inhibition, nearly reaching the level observed upon BRCA1 depletion, which served as a positive control (**Figure 7B**). Additionally, we employed the EJ5 NHEJ HeLa reporter system, in which the disrupted GFP is reactivated by the NHEJ process. Again, a significant reduction in NHEJ efficiency was observed after INTS6 depletion (wortmannin, a DNA-PK inhibitor was used as the positive control) (**Supplementary Figure S10A**). Comet assay results, visualizing broken DNA ends in wildtype and INTS6-depleted cells (**Figure 7C and Supplementary Figure S10B**, siRAD51 was used as positive control), showed that at 0.5 hours post-IR, broken DNA ends were evident in all samples. However, at 24 hours post-IR, the broken ends in control cells had mostly been repaired, whilst siINTS6 cells exhibited a comparable number of broken ends to siRAD51 cells. Collectively, these data indicate that INTS6 is indispensable for efficient DNA repair.

**Figure 7.**
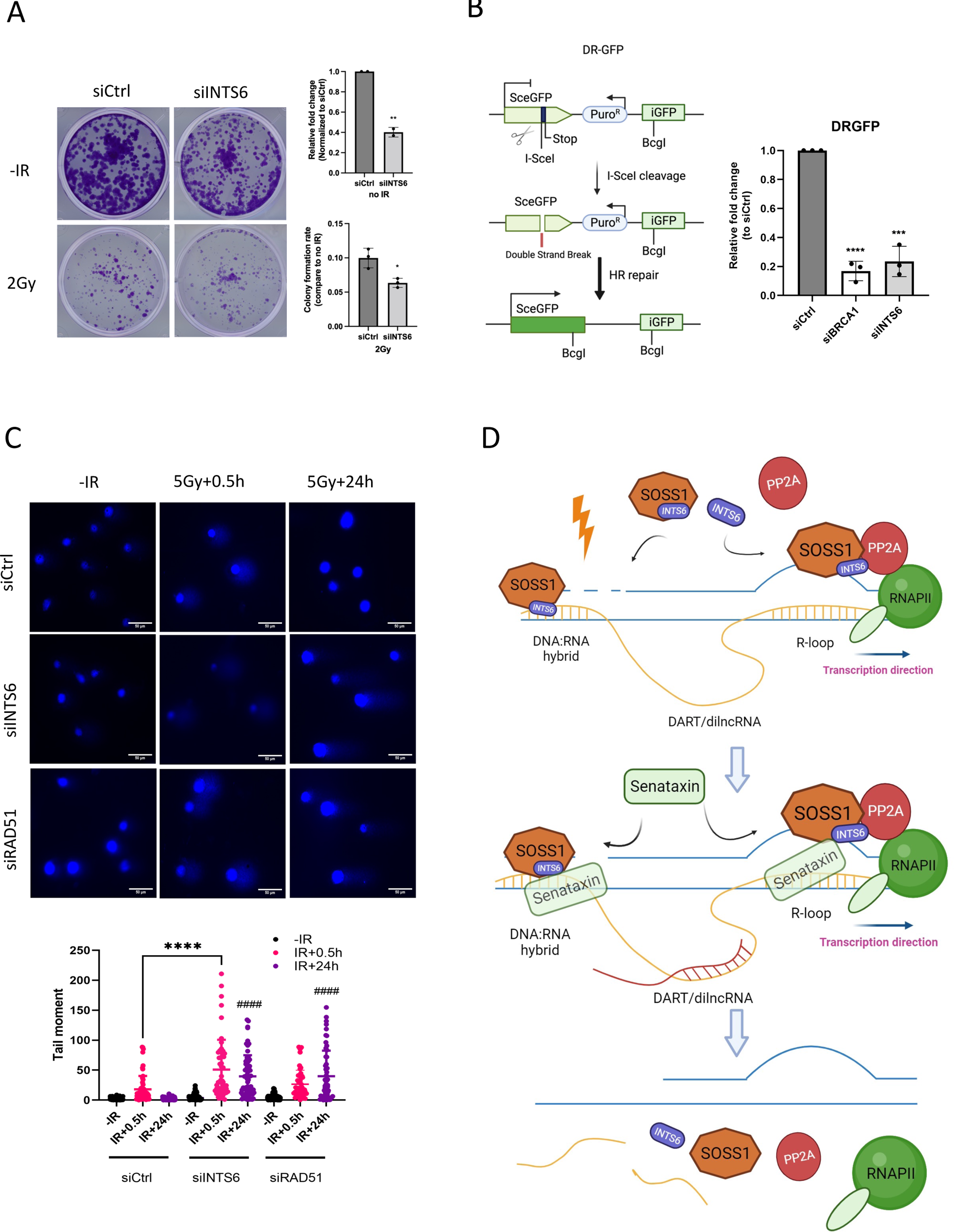
INTS6 is required for efficient DNA damage repair. **A)** Left: representative images of the clonogenic assay in control and INTS6 knockdown cells. The cells were stained and counted after 10 days of growing. Right: quantification of left. * p ≤0.05, **p ≤ 0.01. **B)** Left: Drawing of DR-GFP HR reporter strategy. Right: Bar chart shows the efficiency of HR repair in DR-GFP HeLa reporter cells, as measured by FACS. BRCA1 knockdown was used as the positive control. **C)** Left: Representative images of comet assay in siCtrl, siINTS6 and siRAD51 cells. siRAD51 cells work as the positive control. IR=5Gy. Error bar = 50 μm. ****p ≤ 0.0001 represents the comparison between siCtrl and siINTS6, ^####^p ≤ 0.0001 represents the comparison between 24h and 0.5h. **D)** Proposed model: INTS6 as part of tetrameric SOSS1 complex binds to DNA:RNA hybrids at DSBs and recruits PP2A to dephosphorylate RNAPII. Depletion of INTS6 results in increased levels of DARTs and DNA:RNA hybrids. INTS6 interacts with Senataxin and is required for its recruitment to damaged sites. Senataxin in turn resolves DNA:RNA hybrids at DSBs facilitating their INTS6-dependent autoregulation.

Based on our findings, we proposed that INTS6, as part of the tetrameric SOSS1 complex, binds to DNA:RNA hybrids at DSBs and recruits PP2A to dephosphorylate RNAPII. Depletion of INTS6 results in increased levels of DARTs and DNA:RNA hybrids. INTS6 interacts with SETX and is required for its recruitment to damaged sites. SETX, in turn, resolves DNA:RNA hybrids at DSBs, facilitating their INTS6-dependent autoregulation.

## Discussion

Transcription by RNA polymerase II (RNAPII) is a highly regulated process essential for cellular function. The Integrator complex, evolutionarily conserved across metazoans and comprising 16 subunits(29), exerts regulatory control over the fate of numerous nascent RNAs transcribed by RNAPII. With an inherent RNA endonuclease activity, Integrator contributes to the biogenesis of small nuclear RNAs and enhancer RNAs. Significantly, the Integrator complex is instrumental in initiating premature transcription termination at various protein-coding genes, resulting in the attenuation of gene expression. Consequently, the Integrator complex plays a pivotal role in shaping the transcriptome, ensuring the robust inducibility of genes when necessary (29,51,63). When coupled with PP2A Phosphatase, the Integrator complex forms the Integrator-PP2A complex, regulating the dephosphorylation of active RNAPII (38).

Even though the molecular structure of the Integrator has been solved by Cryo-EM(38,64), showing the exact interaction architecture of the subunits within the Integrator, there is evidence for different interacting patterns between various subunits, suggesting that they might co-exist and function outside of the Integrator complex. For instance, INTS3 and INTS6 are parts of the different modules not in close proximity with each other within the Integrator complex, yet they can interact as part of the SOSS1 complexes(41). The SOSS1 complex, a multiprotein complex, is crucial in the DNA damage response, functioning downstream of the MRN complex to promote DNA repair and the G2/M checkpoint. Essential for efficient homologous recombination-dependent repair of double-strand breaks (DSBs) and ATM-dependent signalling pathways, the SOSS1 complex acts as a sensor of single-stranded DNA, particularly binding to polypyrimidines(65). Moreover, the trimeric SOSS1 complex promotes phase separation at DSBs(39).

This study reveals that in response to DNA damage, INTS6 binds to the trimeric SOSS1 to form the tetrameric SOSS1 complex. The tetrameric SOSS1 then recruits PP2A to DSBs, facilitating the dephosphorylation of RNAPII. Transcription at DSBs is crucial for efficient DNA repair, but it eventually needs to be terminated. The termination process, associated with the interaction of RNAPII and the cleavage/polyadenylation complex, relies on the S2P CTD of RNAPII. The release of mRNA and the initiation of a new cycle for subsequent rounds require dephosphorylation of the CTD of RNAPII. Thus, the INTS6-dependent recruitment of PP2A can be seen as a prerequisite for efficient transcription termination at DSBs. Furthermore, this study finds that INTS6 interacts with and recruits SETX to DSBs. SETX, recently identified as a transcription termination factor in mammalian cells, suggests a dual role for INTS6 in the regulation of transcription termination at DSBs (46).

DNA:RNA hybrids and R-loops are recognized as potential threats to genome stability (21,27,28). However, they are closely associated with transcription at DSBs and can serve as binding platforms for numerous DDR factors (11,19,23–26). Our study unveils the ability of INTS6 to bind to DNA:RNA hybrids, a crucial step for its localization to DSBs. Depletion of INTS6 results in a significant increase in nascent transcripts at DSBs, known as damage-induced RNA transcripts (DARTs). As DARTs are produced as single-stranded RNA, they can hybridize with the DNA overhang post-resection, forming DNA:RNA hybrids, and to the exposed single-strand DNA behind pausing RNA polymerase II (RNAPII) to create R-loops. Notably, within 1kb on each side of the DSB, the enriched INTS6-mediated transcripts overlap with DNA:RNA DRIP-seq peaks and SETX ChIP-seq peaks. SETX, a well-known DNA:RNA helicase, resolves DNA:RNA hybrids at DSBs (21). This study uncovers a previously unknown interaction between INTS6 and SETX, shedding light on the SOSS1-dependent resolution of DNA:RNA hybrids and their autoregulation.

Overall, we demonstrate the formation of a tetrameric SOSS1 complex, comprising INTS6 and the trimeric SOSS1, in response to DNA damage. INTS6’s specific binding to DNA:RNA hybrids plays a pivotal role in recruiting PP2A to DSBs, facilitating the dephosphorylation of RNAPII. Depletion of INTS6 leads to the accumulation of damage-induced RNA transcripts and the stabilization of DNA:RNA hybrids at DSB sites. Additionally, INTS6 interacts with and mediates the recruitment of SETX to DSBs, facilitating the resolution of DNA:RNA hybrids. These findings underscore the critical role of the SOSS1 complex in autoregulating DNA:RNA dynamics and promoting effective DNA repair.

## Data availability

Data reported in this paper can be shared by the lead contact upon request. Mass spectrometry proteomics data have been deposited to the ProteomeXchange Consortium via PRIDE66.

ChrRNA-seq data have been deposited to GEO and can be accessed under GSE246729 with private token: mlwbwamidfgvpol

Any additional information required to re-analyze the data reported in this work paper is available from the lead contact upon request.

## Acknowledgements

This work was supported by the Senior Research Fellowship by Cancer Research UK [grant number BVR01170], EPA Trust Fund [BVR01670], and Lee Placito Fund awarded to M.G., Junior Star Grant from the Grant Agency of the Czech Republic (21-10464M) awarded to M.S. Additional funding included the European Research Council (ERC) under the European Union’s Horizon 2020 research and innovation programme (grant agreement No. 649030 to R.S.), which supported initial experiments. R.S. was additionally supported by the grant CZ.02.01.01/00/22_008/0004575 RNA for therapy, funded by Ministry of Education, Youth, and Sports of the Czech Republic. This work was also supported by funding from University of Miami Miller School of Medicine, Sylvester Comprehensive Cancer Center and grants R01GM078455 from the National Institute of Health to R.Sh.

We are grateful to Prof. Fumiko Esashi (University of Oxford) for providing us with the AsiSI-ER U2OS and wild-type cells which were originally from Legube Lab (CNRS – University of Toulouse, France); and for providing us with homemade RAD51 antibody; Dr Sue Tan-Wang from Prof. Nicholas Proudfoot’s lab (University of Oxford) for sharing the pRNH1-GFP, pRNH1D210N-GFP and pRNH1WKKD-GFP plasmids.

## Author contributions

Q. L. performed most of the experiments and prepared initial draft of the manuscript. K.S. and V.H. purified proteins and performed EMSA, MTS, and pull-down experiments with the assistance. K.A. performed all of the bioinformatics analysis. R.S. supervised and funded work at Ceitec. M.S. designed, supervised, and funded work at Ceitec. S.D., F. B. and M.G.V. performed the affinity purification and analyses of affinity fractions by silver staining and western blot analyses. Mass spectrometry was performed at Wistar institute. R.Sh. supervised and funded work at Miami. M.G. designed and supervised the project and wrote the manuscript with R. Sh. and M.S. assistance. M.G. also assisted with microscopy. All the authors reviewed and approved the final version of the manuscript.

---

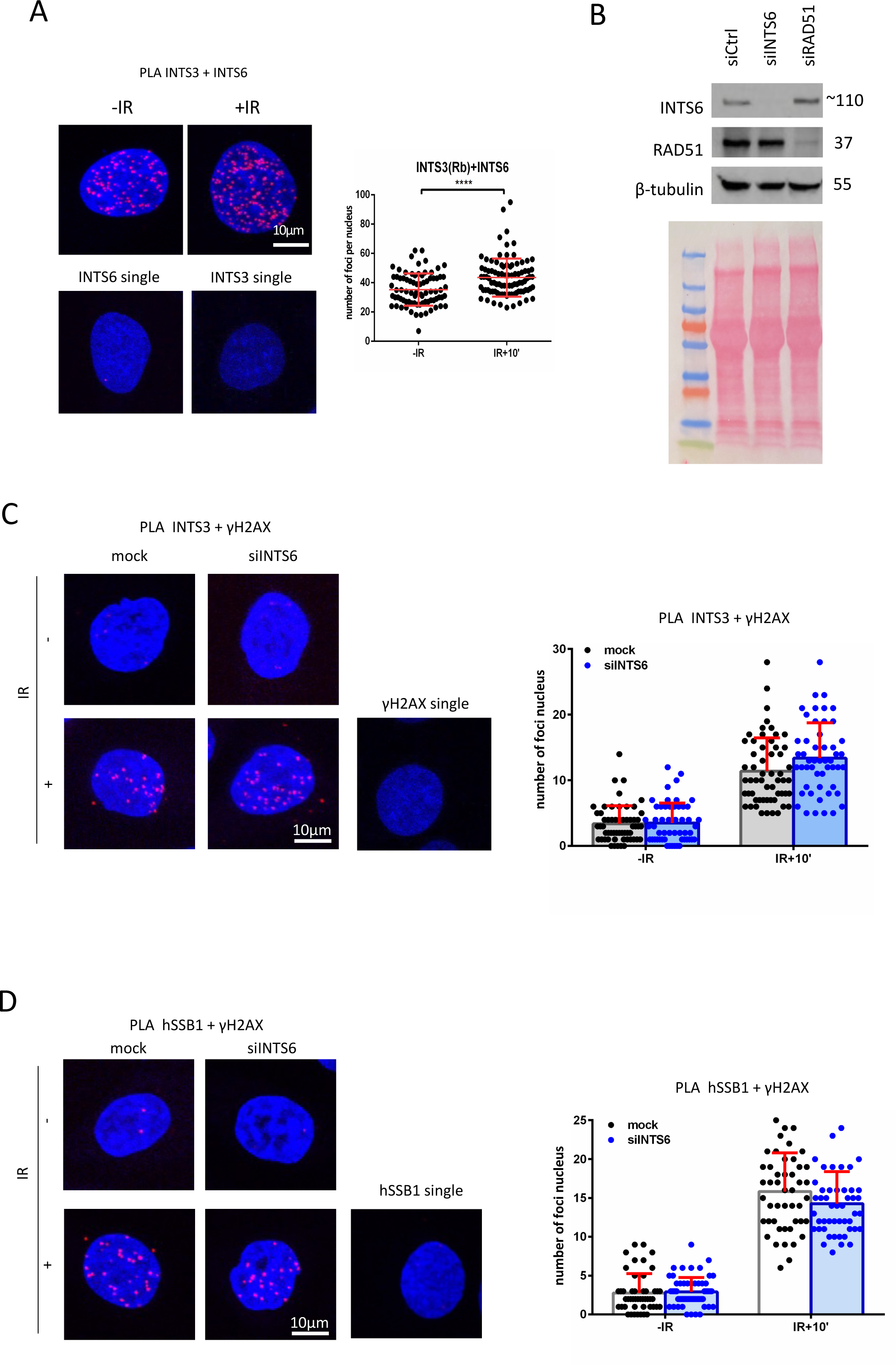
Supplementary Figure 1

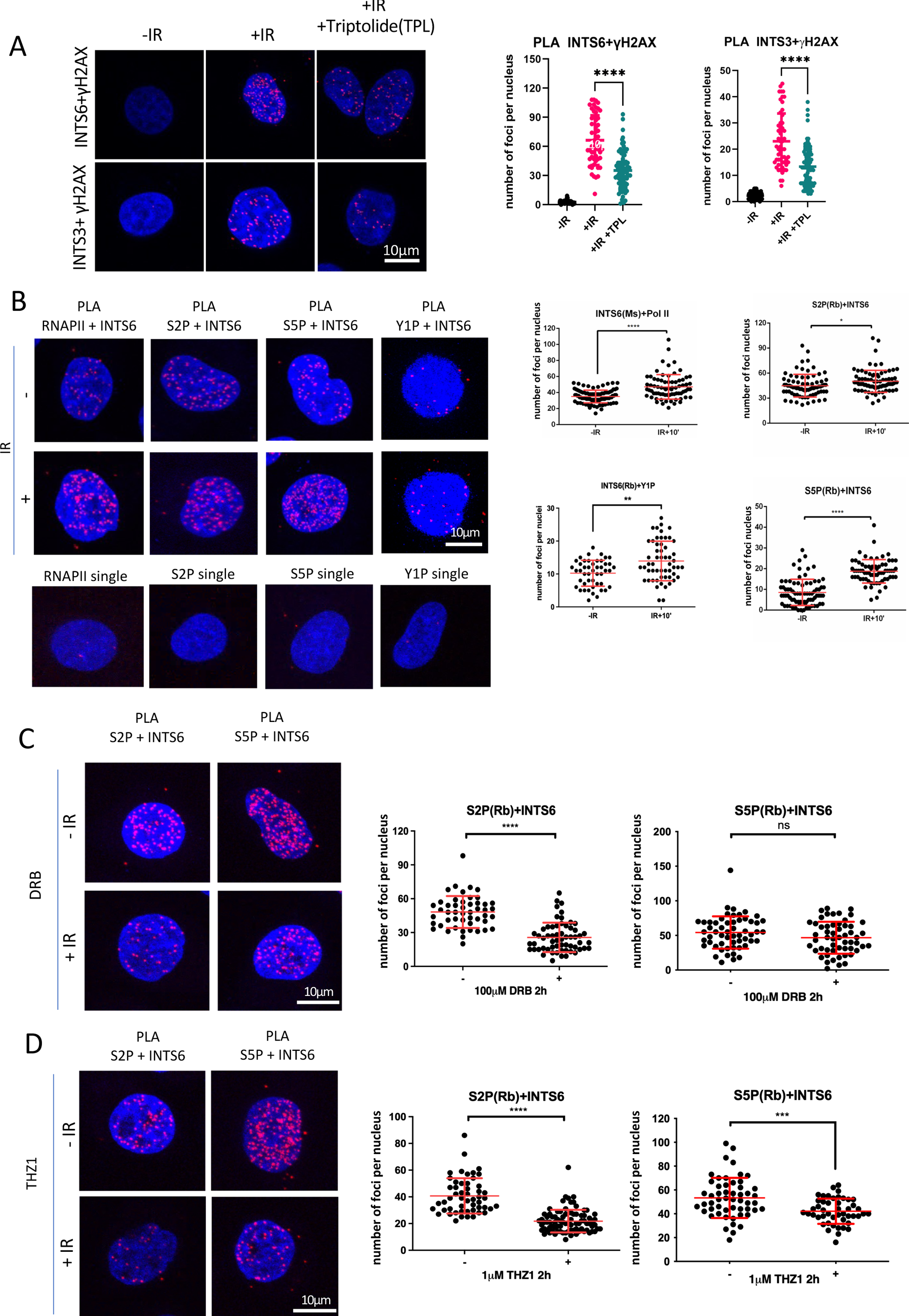
Supplementary Figure 2

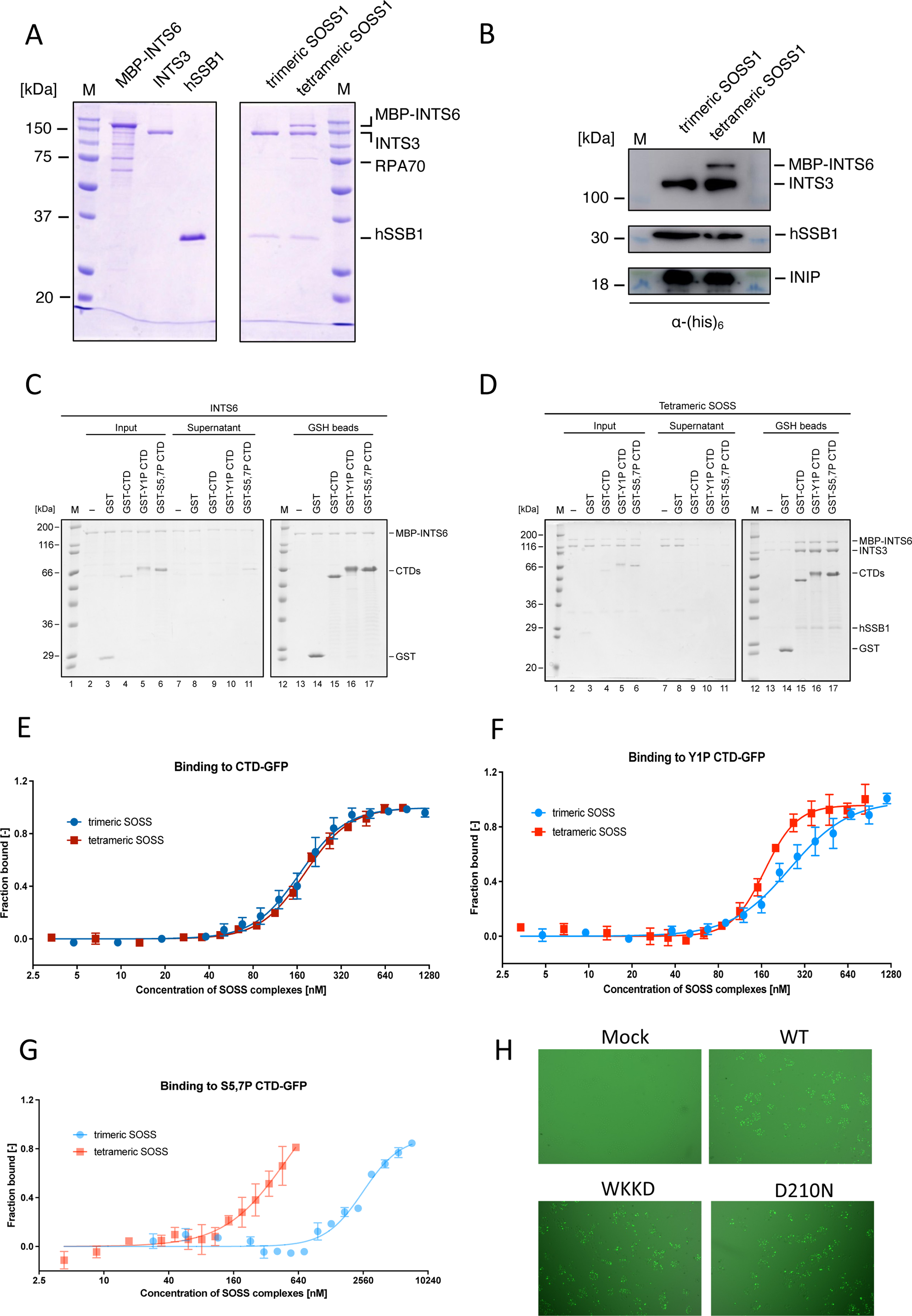
Supplementary Figure 3

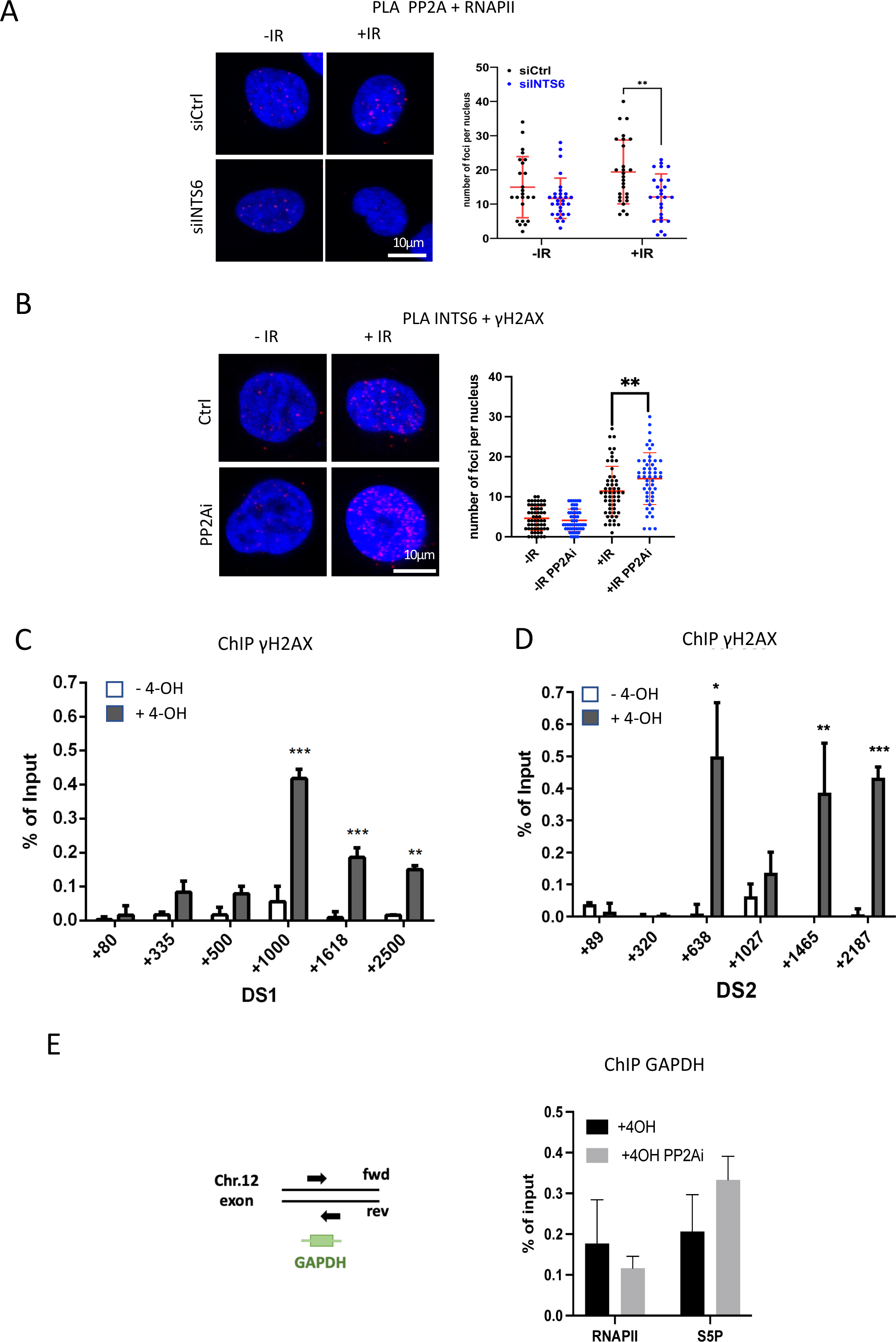
Supplementary Figure 4

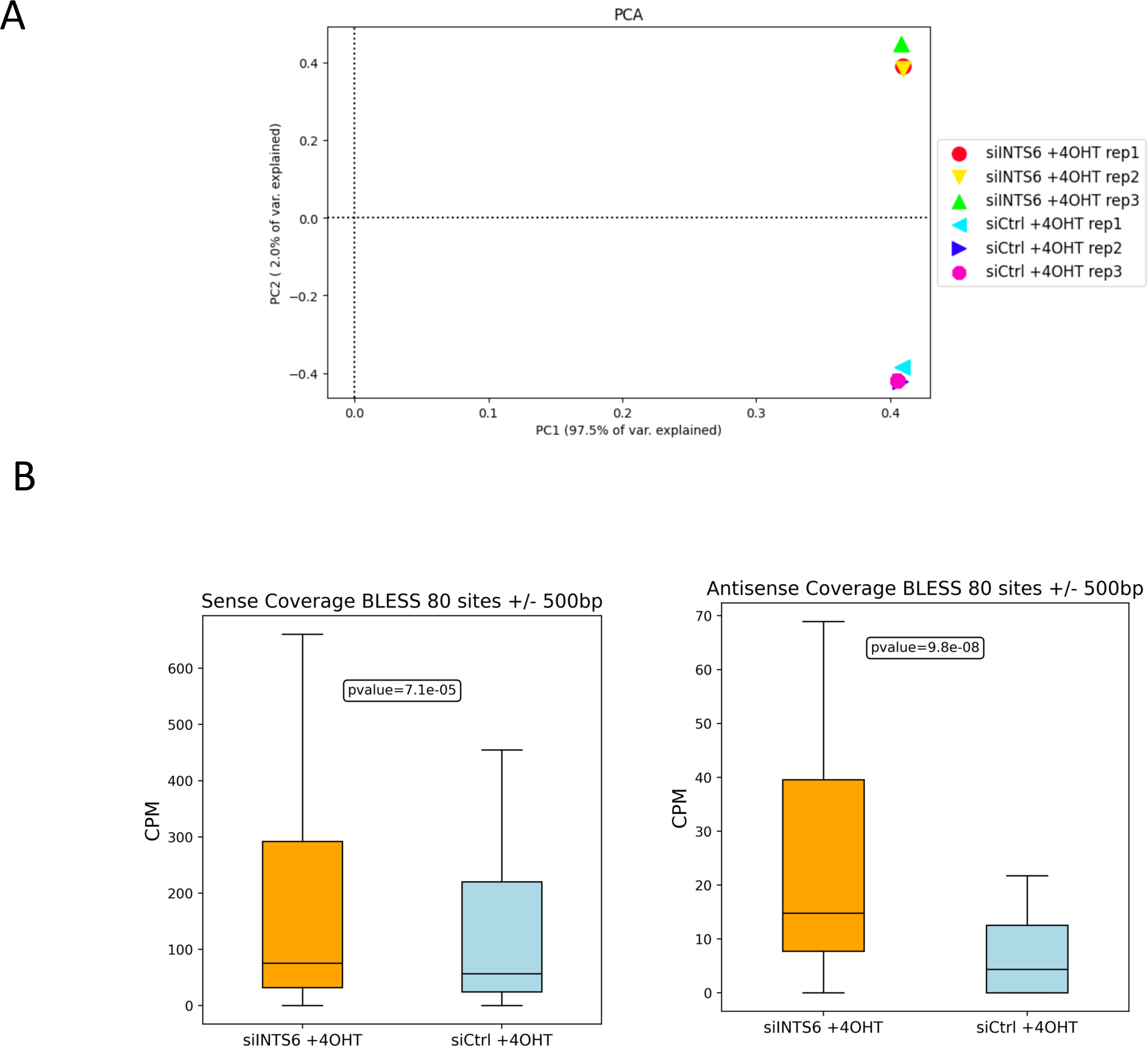
Supplementary Figure 5

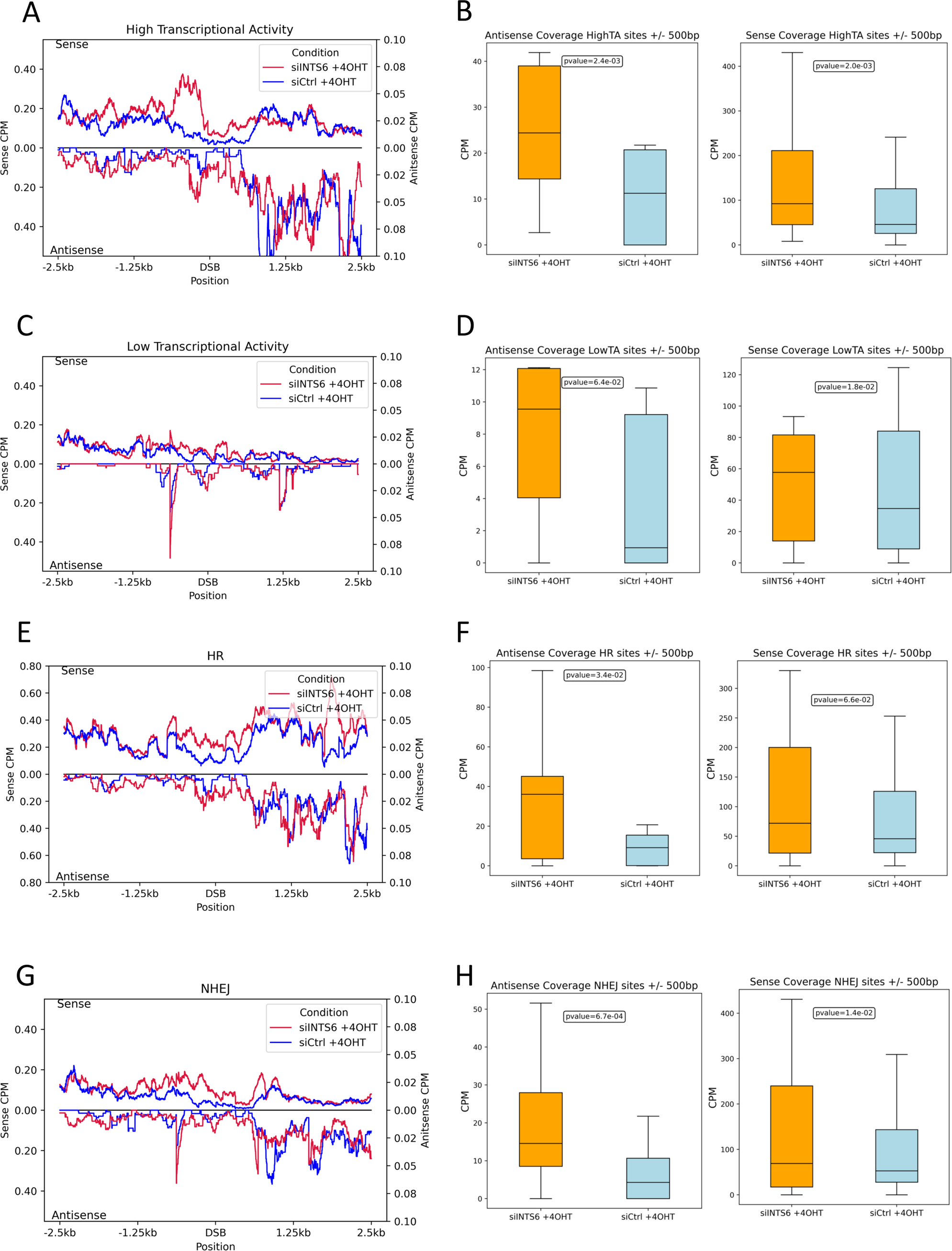
Supplementary Figure 6

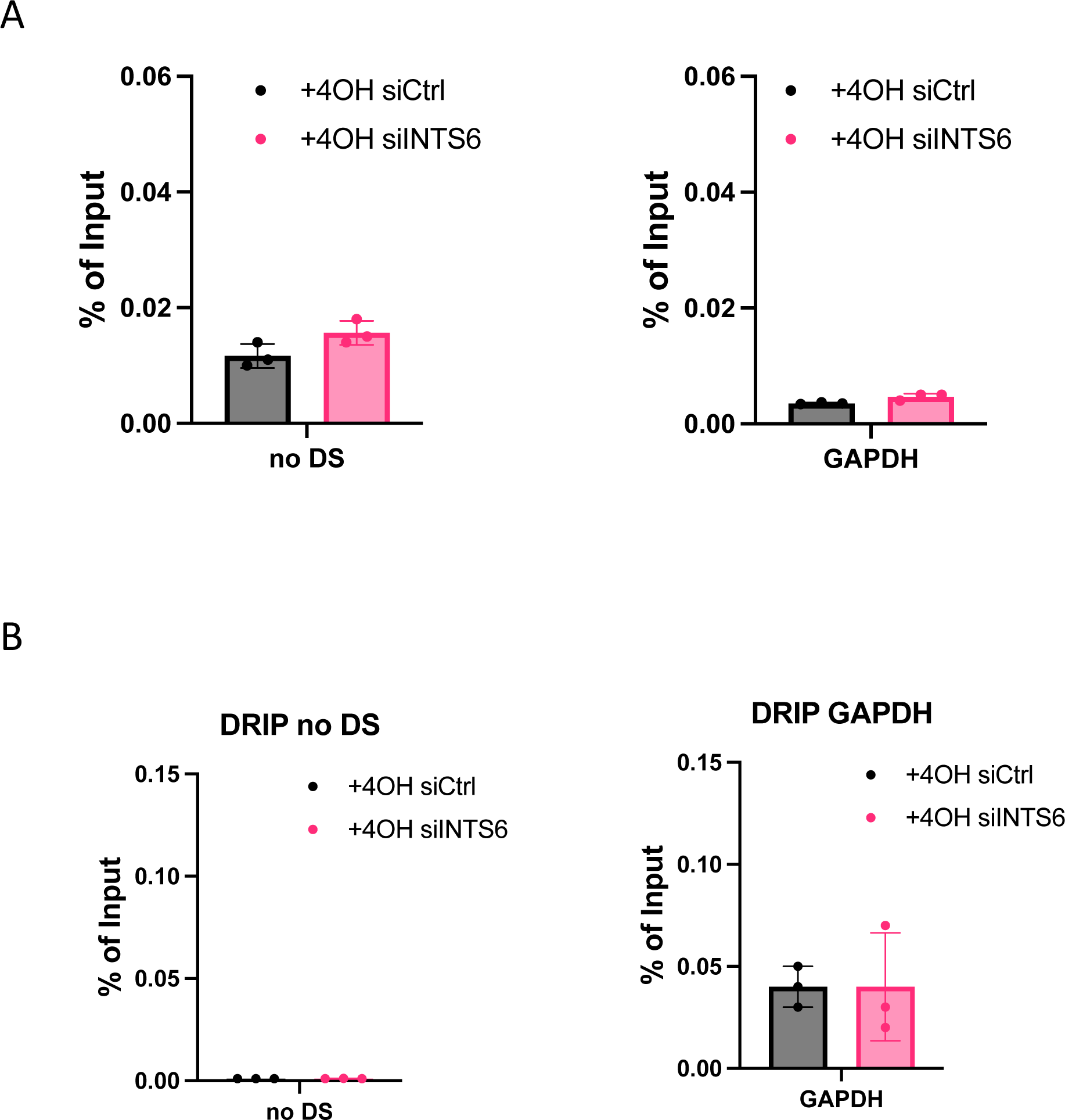
Supplementary Figure 7

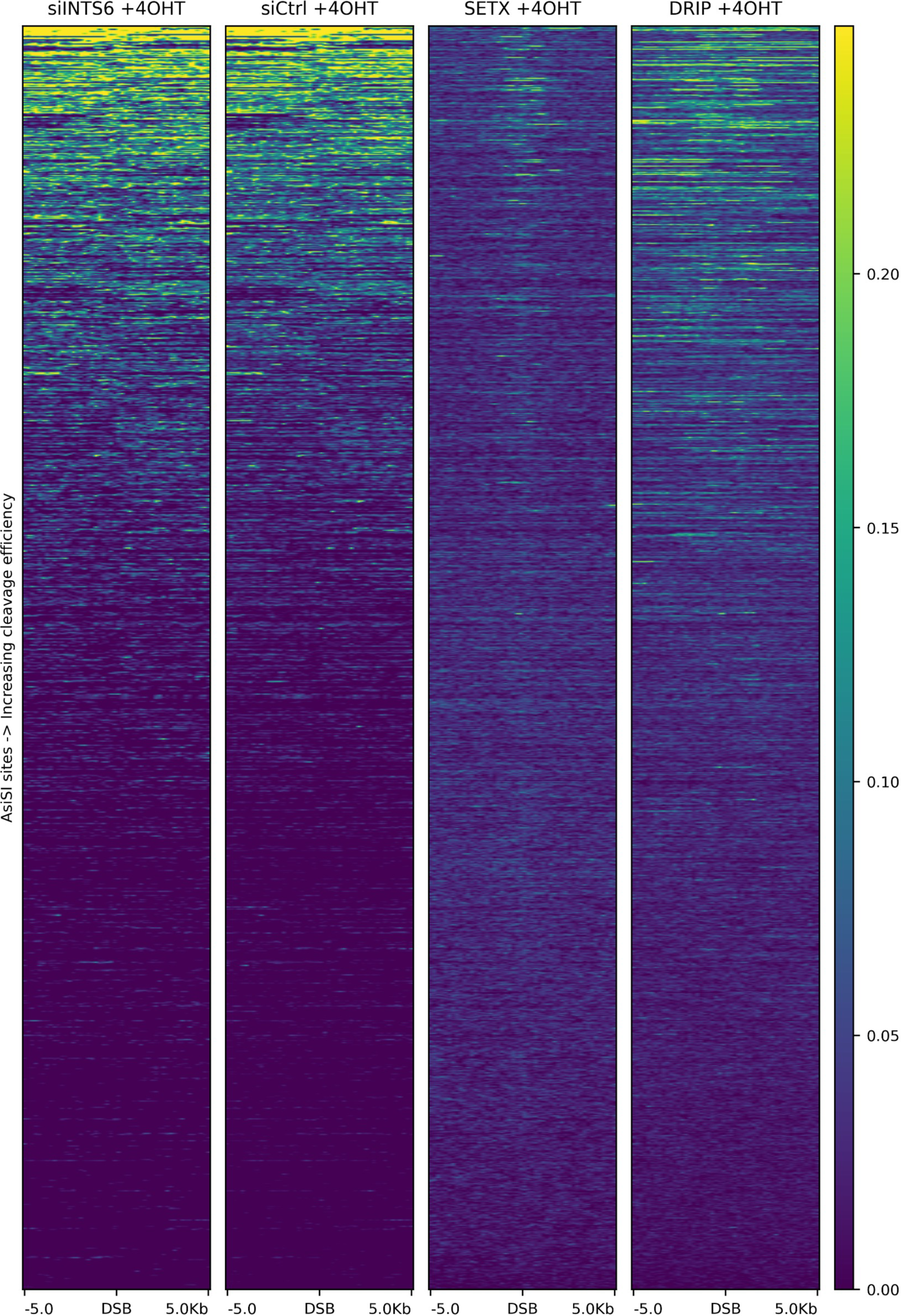
Supplementary Figure 8

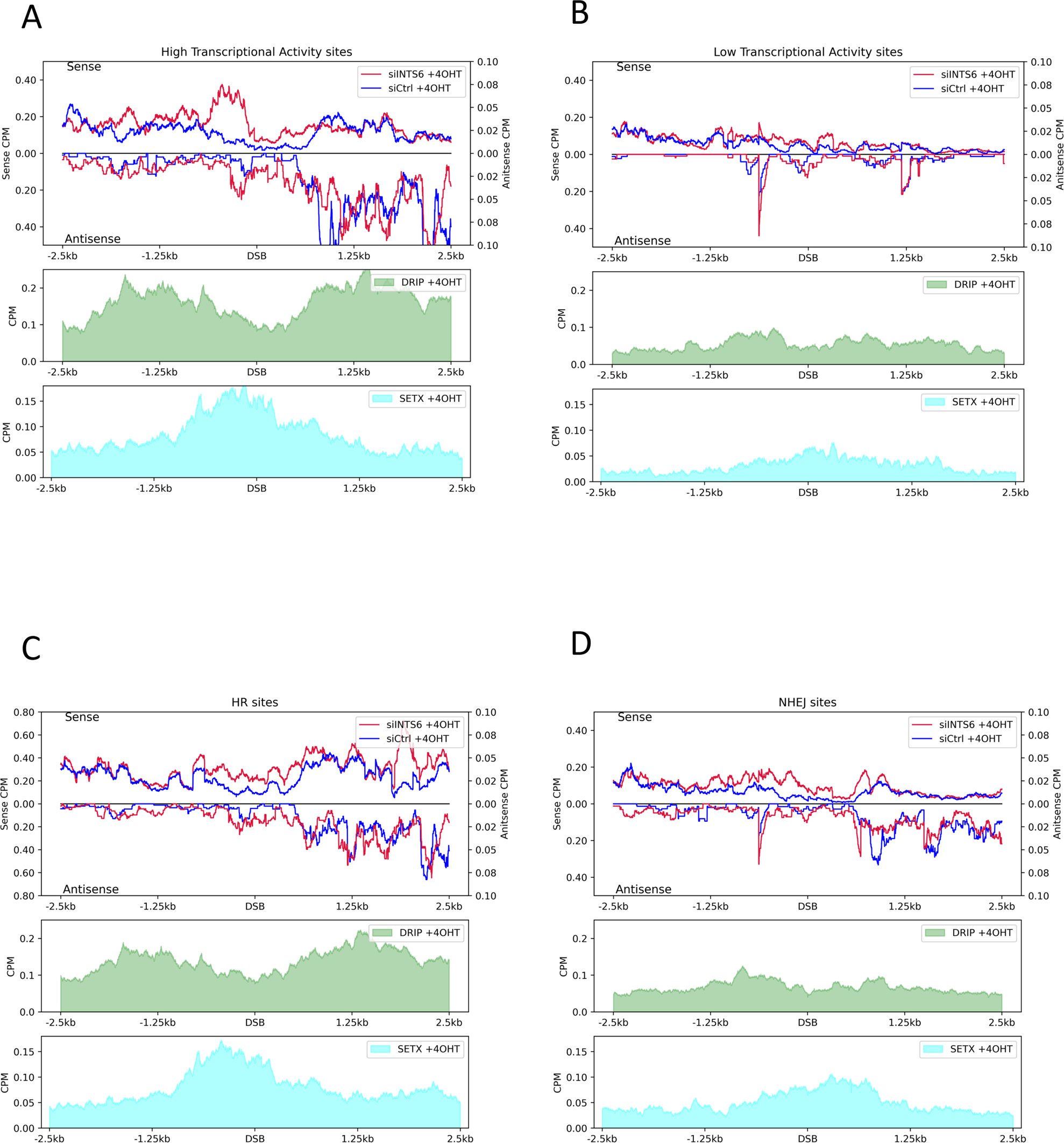
Supplementary Figure 9

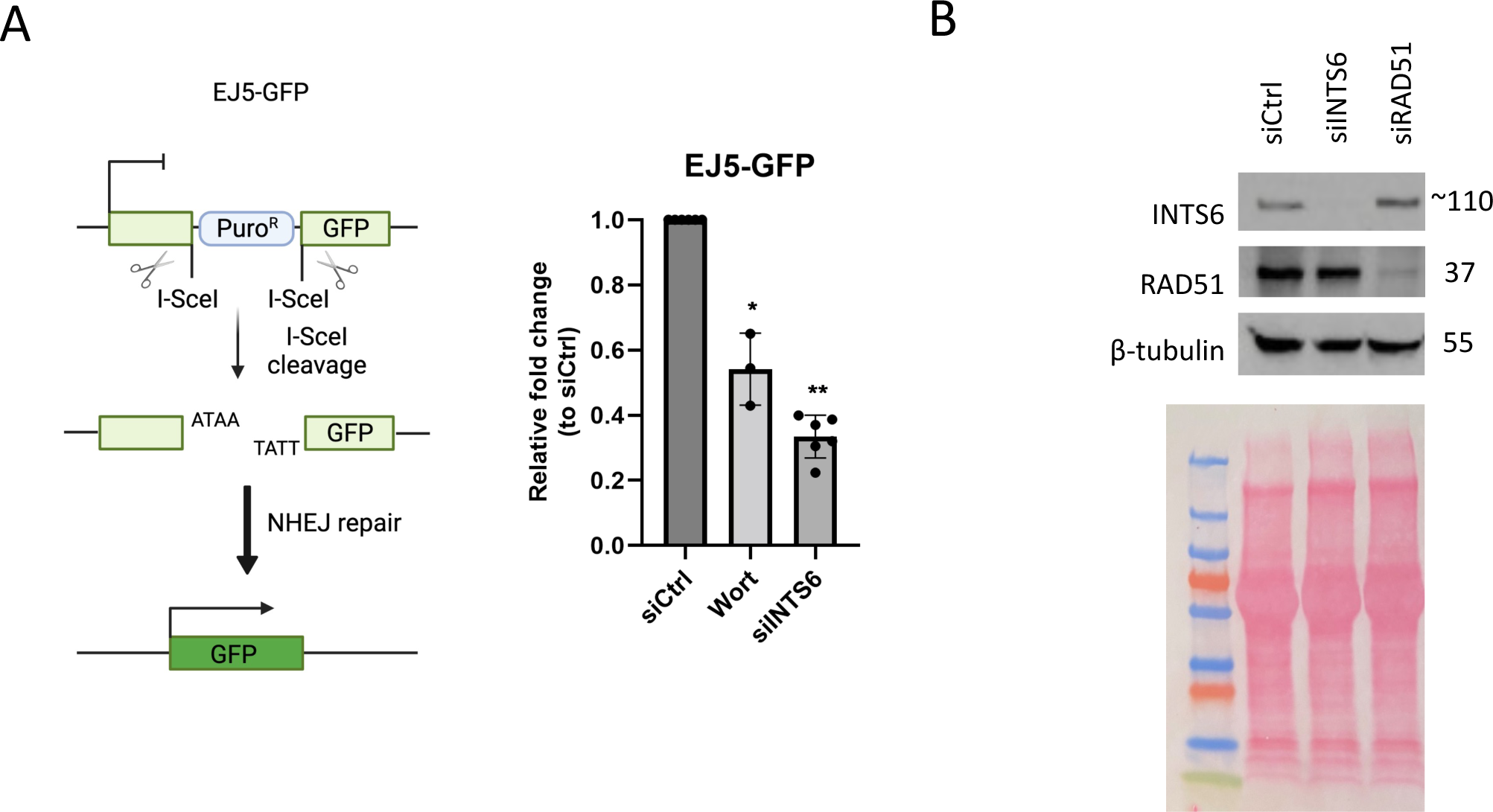
Supplementary Figure 10

