## Supplementary Figure legends for "Tetrameric INTS6-SOSS1 complex facilitates DNA:RNA hybrid autoregulation at double-strand breaks"

**Supplementary Figure legend**

**Figure S1. Tetrameric SOSS1 forms upon DNA damage**

**A)** PLA of INTS3 and INTS6 in cells with or without IR. IR=10Gy. Left: representative confocal microscopy images; Right: quantification of top, error bar = mean ± SD, significance was determined using non-parametric Mann-Whitney test. ****$p \leq 0.0001$. Scale bar =10μm. Single antibodies were used as a negative control.

**B)** Representative images of western blot showing the efficiency of INTS6 knockdown.

**C-D)** PLA of INTS3 (C) or hSSB1 (D) and γH2AX in mock or INTS6 knockdown cells with or without IR. IR=10Gy. Left: representative confocal microscopy images; right: quantification of left, error bar = mean ± SD, significance was determined using non-parametric Mann-Whitney test. Scale bar =10μm. Single antibodies were used as a negative control.

**Figure S2. Tetrameric SOSS1 binds to phosphorylated CTD of RNAPII in response to DNA damage**

**A)** PLA of INTS3 or INTS6 and γH2AX in cells with or without IR in the presence or absence of triptolide (20μM, 1h). IR=10Gy. Left: representative confocal microscopy images; right: quantification of left, error bar = mean ± SD, significance was determined using non-parametric Mann-Whitney test. ****$p \leq 0.0001$. Scale bar =10μm.

**B)** PLA of INTS6 and RNAPII or S2P or S5P or Y1P with or without IR. IR=10Gy. Top: representative confocal microscopy images; bottom: quantification of top, error bar = mean ±SD, significance was determined using non-parametric Mann-Whitney test. *$p \leq 0.05$, **$p \leq 0.01$****$p \leq 0.0001$. Scale bar =10μm. Single antibodies were used as a negative control.

**C)** PLA of S2P or S5P and INTS6 with or without 100μM DRB treatment for 2h. IR=10Gy. Left: representative confocal microscopy images; right: quantification of left, error bar = mean ± SD, significance was determined using non-parametric Mann-Whitney test. ****$p \leq 0.0001$. Scale bar =10μm.

**D)** PLA of S2P or S5P and INTS6 in cells with or without 1μM THZ1 treatment for 2h. IR=10Gy. Left: representative confocal microscopy images; right: quantification of left, error bar = mean ± SD, significance was determined using non-parametric Mann-Whitney test. ***$p \leq 0.001$, ****$p \leq 0.0001$. Scale bar =10μm.

**Figure S3. Tetrameric SOSS1 binds to phosphorylated CTD of RNAPII with higher affinity than trimeric SOSS1**

**A)** A representative 15% SDS-PAGE gel depicting purified components of SOSS1 complexes (INIP is not visible due to its size).

**B)** 18% SDS-PAGE followed by western blot probing for components of trimeric or tetrameric SOSS1 complexes.

**C)** A representative SDS-PAGE gel depicting *in vitro* pull-down assay of INTS6 with GST-tagged CTD, GST-tagged CTD modified on tyrosine 1 (Y1P), or GST-tagged CTD modified on serine 5 and 7 (S5,7P).

**D)** A representative SDS-PAGE gel depicting *in vitro* pull-down assay of tetrameric SOSS1 complex with GST-CTD, GST-Y1P-CTD or GST-S5,7P-CTD.

**E-G)** Microscale-thermophoresis (MST) binding curves of trimeric and tetrameric SOSS1 complexes with unmodified CTD-GFP (E), Y1P-CTD-GFP (F) or S5,7P-CTD-GFP (G). Measured in triplicates and the lines represent the Hill fit.

**H)** Confocal microscopy images showing GFP expression of transiently transfected RNAseH1 $^{wt\text{-}GFP}$ or RNAseH1 $^{WKKD\text{-}GFP}$ (binding and catalytic) or RNAseH1 $^{D210N\text{-}GFP}$ (catalytic) mutants. Mock is used as a negative control.

**Figure S4. The recruitment of INTS6 to DSBs after PP2Ai and the validation of *Asi*SI-ER cut**

**A)** PLA of PP2A and RNAPII with or without IR in the control or INTS6 knockdown cells. IR=10Gy. Left: representative confocal microscopy images; right: quantification of left, error bar = mean ± SD, significance was determined using non-parametric Mann-Whitney test. $^{**}p \leq 0.01$. Scale bar =10μm.

**B)** PLA of INTS6 and γH2AX with or without IR in the presence or absence of PP2A inhibitor (LB-100, 2.5μM, 2h). IR=10Gy. Left: representative confocal microscopy images; right: quantification of left, error bar = mean ± SD, significance was determined using non-parametric Mann-Whitney test. $^{**}p \leq 0.01$. Scale bar =10μm.

**C)** Bar chart showing γH2AX ChIP signals around DS1 in the absence or presence of 4-OHT. n=3, significance was determined by student t-test, $^{***}p \leq 0.001$, $^{**}p \leq 0.01$

**D)** Bar chart showing γH2AX ChIP signals around DS2 in the absence or presence of 4-OHT. n=3, significance was determined by student t-test, $^{*}p \leq 0.05$, $^{**}p \leq 0.01$, $^{***}p \leq 0.001$.

**E)** Left: drawing showing position of ChIP probe in GAPDH gene. Right: Bar chart showing RNAPII and S5P RNAPII ChIP signals at GAPDH gene in the absence of 4-OHT and presence or absence of PP2A inhibitor (LB-100, 2.5µM, 2h).

**Figure S5. ChrRNA-seq quality control**

**A)** PCA plots show a comparison of chrRNA-Seq coverage in the 5kb flanking region of BLESS 80 *Asi*SI sites between sample replicates in INTS6 knockdown and control conditions.

**B)** Box plot shows chrRNA-Seq sense (Left) and antisense (Right) coverage in +/- 500 bp flank of BLESS 80 *Asi*SI sites in INTS6 knockdown and control after 4-OH treatment. Wilcoxon 2 sample test is used with medians test. Error bar = mean ± SD.

**Figure S6. Coverage of chrRNA-Seq at DSBs with certain features**

**A)** Metagene plots showing chrRNA-Seq sense and antisense coverage in INTS6 knockdown and control cells with damage induction (4-OHT) around 2.5kb flank region of highly transcriptionally active *Asi*SI sites (n=20).

**B)** Box plot shows chrRNA Seq antisense (Left) and sense (Right) coverage in +/- 500 bp flank of highly transcriptionally active *Asi*SI sites (n=20).

**C)**, As in A) for transcriptionally inactive *Asi*SI sites (n=20)

**D)** As in B) for transcriptionally inactive *Asi*SI sites (n=20).

**E)** As in A) for HR prone *Asi*SI sites (n=30)

**F)** As in B) for HR prone *Asi*SI sites (n=30)

**G)** As in A) for NHEJ prone *Asi*SI sites (n=30)

**H)** As in B) for NHEJ prone *Asi*SI sites (n=30)

**Figure S7. SETX ChIP and DRIP control regions**

**A)** Bar chart showing SETX ChIP signals over no DSB locus (chr22:23141639-23141780) and GAPDH, in the presence or absence of INTS6. N=3. Error bar = mean ± SD.

**B)** Bar chart showing SETX DRIP signals over no DSB locus (chr22:23141639-23141780) and GAPDH, in the presence or absence of INTS6. N=3. Error bar = mean ± SD.

**Figure S8. Heatmaps on BLESS 80 *Asi*SI sites**

Heatmaps show the siINTS6 chrRNA-Seq coverage, Ctrl chrRNA-Seq coverage, SETX ChIP-Seq coverage and S9.6 DRIP-Seq coverage across all annotated *Asi*SI sites. The reference genome is human hg19.

**Figure S9. Comparison of nascent RNA transcripts, DNA:RNA hybrids and SETX at different categories of *Asi*SI sites**

**A)** Metagene plots showing DRIP and SETX ChIP coverage upon damage induction along with sense and antisense coverage of chrRNA-Seq in siINTS6 and control with damage induction around 2.5kb flank region of highly transcriptionally active *Asi*SI sites (n=20). The reference genome is human hg19.

**B)** Metagene plots showing DRIP and SETX ChIP coverage upon damage induction along with sense and antisense coverage of chrRNA-Seq in siINTS6 and control with damage induction around 2.5kb flank region of transcriptionally inactive *Asi*SI sites (n=20). The reference genome is human hg19.

**C)** Metagene plots showing DRIP and SETX ChIP coverage upon damage induction along with sense and antisense coverage of chrRNA-Seq in siINTS6 and control with damage induction around 2.5kb flank region of HR-prone *Asi*SI sites (n=30). The reference genome is human hg19.

**D)** Metagene plots showing DRIP and SETX ChIP coverage upon damage induction along with sense and antisense coverage of chrRNA-Seq in siINTS6 and control with damage induction around 2.5kb flank region of NHEJ-prone *Asi*SI sites (n=30). The reference genome is human hg19.

**Figure S10. Western blot shows the knockdown efficiency**

**A)** Left: Drawing of EJ5-GFP NHEJ reporter strategy. Right: Bar chart showing the efficiency of NHEJ repair in EJ5 HeLa reporter cells, as measured by FACS. Wortmannin (DNA-PK inhibitor) was used as the positive control.

**B)** Western blot shows the knockdown efficiency of INTS6 and RAD51 in HeLa cells.
