## Supplementary Tables for "Tetrameric INTS6-SOSS1 complex facilitates DNA:RNA hybrid autoregulation at double-strand breaks"

**Table S1. List of oligonucleotides used in this study for *in vitro* work.**

| Name | Sequence 5'-3' Supplementary table | Use |
|---|---|---|
| pR209 | TACTTCCAATCCAATCGATGACGACGGAGACCTTTG | Amplification of INTS6 |
| pR210 | TTATCCACTTCCAATGTTATTACTAATTGCTATTAATATGGTTGATC | Amplification of INTS6 |
| pR211 | GAACAGTTGACAGGTGTGCC | Sequencing of INTS6 |
| pR212 | TCCAGTCCTTCTTCCCCTCT | Sequencing of INTS6 |
| pR213 | CAAATGGGGAACTACCAGGA | Sequencing of INTS6 |
| Part of the following substrates: RNA:DNA-hybrid, R-loop | | |
| pR219 | GGGTGAACCTGCAGGTGGGCGGCTGCTCATCGTAGGTTAGTTGGTAGAATTCGGCAGCGTC | EMSA experiments |
| Part of the following substrate: R-loop substrate for EMSA experiments | | |
| pR223 | AAAGAUGUCCUAGCAAGGCAC | EMSA experiments |
| Part of the following substrates: RNA:DNA-hybrid, R-loop | | |
| pR630 | [Cy3]-GTGCCTTGCTAGGACATCTTT | EMSA experiments |

**Table S2. List of oligonucleotides used in this study for *in vivo* work.**

| Primer Name | Primer Sequence (5'-3') |
|---|---|
| DS1-335 fwd | GAATCGGATGTATGCGACTGATC |
| DS1-335 rev | TTCCAAAGTTATTCCAACCCGAT |
| DS1-500 fwd | CCTGGATATGAGTTTGATCAGC |
| DS1-500 rev | CTCTCCTTTCGCTGACACTG |
| DS1-1000 fwd | AGGAATTGACTGCGGTGTTC |
| DS1-1000 rev | GGGGAGGAGGAAAGGTGTAG |
| DS1-1618 fwd | TGAGGAGGTGACATTAGAACTCAGA |
| DS1-1618 rev | AGGACTCACTTACACGGCCTTT |
| DS1-2500 fwd | GCCATAACAGAGGGTGGAAA |
| DS1-2500 rev | AACTTTAGGATGGGGCTGCT |
| DS2-320 fwd | CTAGGTCTGGCTCCTCCTGA |
| DS2-320 rev | CTCCCTGAACCGCCTAGAAC |

| | |
|---|---|
| DS2-638 fwd | GCTGCCTGAGATGCCTGTAA |
| DS2-638 rev | ATCCAGACAGGCTCCTCCTC |
| DS2-1027 fwd | CCCCAGCTCCTTAACACAGA |
| DS2-1027 rev | CATGGTGCAGAAGGTGCATT |
| DS2-1465 fwd | AGCCCAGTGGCACAGGAATA |
| DS2-1465 rev | CCTTCAGGGGTGACACATCAG |
| DS2-2187 fwd | AGGAATCCACCTATCCGCCT |
| DS2-2187 rev | GGCTAACAGACTTCCAGGCA |
| GAPDH fwd | AACCTGCCAAATATGATGAC |
| GAPDH rev | AGGAAATGAGCTTGACAAAG |
| no DSB-fwd | ATTGGGTATCTGCGTCTAGTGAGG |
| no DSB-rev | GACTCAATTACATCCCTGCAGCT |

**Table S3. List of oligonucleotides used in this study for Gibson cloning.**

| Primer Name | Primer Sequence (5'-3') |
|---|---|
| INTS6_EGFP_fwd | Cagctgttgggctcgcggttatgcccatcttactgttcctg |
| INTS6_EGFP_rev | Ctcacagagcctccacccccattgctattaatatggttgatctgattg |
| BACKBONE_EGFP_fwd | Tcaaccatattaatagcaatgggggtggaggctctgtgag |
| BACKBONE_EGFP_rev | Aggaacagtaagatgggcataaccgcgagcccaacagctg |

**Table S4. List of reagents used in this study.**

| REAGENT or RESOURCE | SOURCE | IDENTIFIER |
|---|---|---|
| Antibodies | | |
| INTS6 | Abcam | ab86369-100ul |
| phospho-Histone H2A.X (Ser139) | Sigma | 05-636 |
| PPP2R1A antibody | Bethyl | A300-962A |
| RNA Polymerase II RPB1-8WG16 | Biolegend | 664912 |
| RNA polymerase II CTD repeat YSPTSPS (phospho S5) | Abcam | ab5131 |
| Anti-RNA polymerase II CTD repeat YSPTSPS antibody | Abcam | ab26721 |
| Rabbit Polyclonal Senataxin Antibody | Nouvs Bio | NBP1-94712 |
| ANTI-DNA-RNA HYBRID, CLONE S9.6 | Sigma | MABE1095 |
| Phospho-gamma-H2AX (Ser139) | Invitrogen | MA533062 |
| INTS3 | bethyl | A302-051A |
| DICE1 (H-6) | Santa Cruz | sc-376524 |
| beta-tubulin | Abcam | ab6046 |
| H3 | Biolegend | 819414 |

| | | |
|---|---|---|
| hSSB1 | Bethyl | A301-938A |
| RNA polymerase II CTD repeat YSPTSPS (phospho S2) | Abcam | ab5095 |
| AbFlex® RNA Pol II CTD phospho Tyr1 antibody (rAb) | Active Motif | 92129 |
| hSSB1 | Lifespan BioSciences | LS-C173584 |
| RAD51 | N/A | FE lab homemade |
| Bacterial and virus strains | | |
| NEB® 5-alpha Competent E. coli (High Efficiency) | New England Biology | C2987H |
| MAX Efficiency$^{TM}$ DH10bac Competent Cells | ThermoFisher | 10361012 |
| Chemicals, peptides, and recombinant proteins | | |
| Triptolide | Enzo life science | BV-1761-1 |
| 5,6-Dichloro-1-beta-D-ribofuranosylbenzimidazole (DRB) | Cayman Chemical | 10010302 |
| THZ1 | Stratech Scientific | A8882-APE-10mM |
| LB-100 | Stratech Scientific | B4846-APE-5mg) |
| (Z)-4-hydroxy Tamoxifen (4OH) | Cayman Chemical | 14854-1mg-CAY |
| Critical commercial assays | | |
| Duolink® In Situ Red Starter Kit Mouse/Rabbit | Sigma | DUO92101-1KT |
| Experimental models: Cell lines | | |
| HeLa | ATCC | N/A |
| DRGFP HeLa | MG lab | N/A |
| EJ5 HeLa | MG lab | N/A |
| U2OS | GL lab | N/A |
| U2OS AsiSI-ER | GL lab | N/A |
| Oligonucleotides | | |
| siControl (ON-TARGETplus, Dharmacon SMARTpool) | Dharmacon | D-001810-03-05 |
| siBRCA1(ON-TARGETplus, Dharmacon SMARTpool) | Dharmacon | J-003461-09-0005 |
| siINTS6(ON-TARGETplus, Dharmacon SMARTpool) | Dharmacon | L-012417-00-0005 |
| siRAD51 #1* | IDT | 5’ GACUGCCAGGAUAAAGCUU 3’ |
| siRAD51 #2* | IDT | 5’ GUGCUGCAGCCUAAUGAGA 3’ |
| *Use both siRAD51 #1 and siRAD51 #2 together to do transient knock down. | | |
| Recombinant DNA: Plasmids | | |
| pCBASceI | (Richardson C et al) | Addgene Plasmid #26477 |
| INTS6-GFP | this study | N/A |
| pRNH1$^{WT}$-GFP | NJP Lab | N/A |
| pRNH1$^{D210N}$-GFP | NJP Lab | N/A |
| pRNH1$^{WKKD}$-GFP | NJP Lab | N/A |
| Software and algorithms | | |

| | | |
|---|---|---|
| GraphPad Prism 9 | GraphPad Software, San Diego, California USA, www.graphpad.com | N/A |
| Fiji | (Schindelinet et al) | N/A |
| CellProfiler | (Carpenter et al) | N/A |
| BioRender | https://www.biorender.com/ | N/A |
